# Dynamic stomach-brain electrical coupling in human sleep

**DOI:** 10.1101/2025.11.13.686572

**Authors:** Akshita A. Rao, Martin Dresler, Ignacio Rebollo, Jamie M. Zeitzer, Sarah F. Schoch, Todd P. Coleman

## Abstract

Sleep involves coordinated interactions between the brain and body, yet whether and how peripheral physiology is organized during human sleep is poorly understood. Because gastric rhythms are strongly modulated by arousal states during wakefulness, we hypothesized that gastric electrical rhythms exhibit structured dynamics aligned with cortical oscillations during sleep. Using simultaneous high-density electroencephalography and electrogastrography, we find that gastric activity persists throughout sleep, increases during non-rapid eye movement (NREM), and exhibits infraslow fluctuations that are amplified during NREM. Moreover, gastric activity is synchronized with slow oscillations and sleep spindles, with gastric power tracking spindle occurrence and cortical infraslow cycles. Stomach-brain coupling predicted next-day memory recall beyond cortical measures, whereas gastric infraslow dynamics predicted subjective sleep quality beyond polysomnographic measures. Together, our findings show that even as the brain disconnects from the external world, it maintains structured communication with peripheral organs, reframing sleep as a coordinated, multi-organ physiological state.

---

Sleep supports restoration, memory consolidation, and homeostatic regulation across species (*1*). While typically considered a state of sensory disengagement (*2*), recent evidence suggests that internal coordination between the brain and periphery continues during sleep (*3*). The autonomic nervous system is a major conduit for this communication through bidirectional interactions between cortical activity and visceral functions such as breathing and digestion (*4*). Sleep stages impose distinct organizational modes on this interaction: non-rapid eye movement (NREM) sleep is characterized by parasympathetic dominance and enhanced brain-body synchronization, whereas rapid eye movement (REM) sleep exhibits cortical desynchronization and pronounced autonomic variability (*5*).

Within NREM sleep, cortical slow oscillations and sleep spindles organize large-scale neural communication (*6*). At a slower timescale, infraslow oscillations (ISOs; *<*0.1 Hz) coordinate activity across cortical, thalamic, and brainstem circuits while modulating arousability (*7*). Comparable infraslow fluctuations occur in peripheral physiology, including heart rate (*8*, *9*), respiration (*9*, *10*), and metabolic signals (*11*). These findings suggest that sleep consists of coordinated central-autonomic dynamics operating over tens of seconds, even as sensory engagement with the external environment is reduced. Importantly, however, the aforementioned peripheral signatures primarily index global autonomic state, and it remains unclear whether organ-specific visceral rhythms are integrated into this infraslow architecture of human sleep.

Among visceral systems, the stomach is a compelling candidate for contributing to brain-body coupling (*12*). Gastric contractions are paced by interstitial cells of Cajal (*13*), producing a slow electrical rhythm (∼0.05 Hz) measurable non-invasively using electrogastrography (EGG) (*12*). This gastric rhythm synchronizes with cortical activity during quiet wakefulness (*14*, *15*), suggesting that visceral oscillations may scaffold large-scale neural organization. More broadly, gut-brain signaling is implicated in a range of neurological and gastrointestinal disorders (*16*), underscoring the importance of understanding how these interactions are organized in fundamental physiological states such as sleep. However, whether gastric rhythms contribute to human sleep physiology remains largely unknown. Prior work of gut-sleep interactions has primarily examined indirect markers such as gastric secretion (*17*) or average EGG changes in gastrointestinal disease groups (*18*). A handful of animal studies suggest that experimentally evoked and naturally occurring gastrointestinal signals can reach and modulate cortical activity during sleep (*19*, *20*), but evidence of stomach-brain coupling during human sleep is lacking.

Interoceptive signals are increasingly recognized as contributors to cognitive and affective processes (*21*, *22*), raising the possibility that visceral dynamics during sleep influence sleep-dependent memory consolidation and sleep quality. During brief arousals, visceral inputs may be arriving at inappropriate phases of sleep and may therefore act as markers of arousal that disrupt consolidation processes. In contrast, coordinated visceral rhythms could provide a scaffold for integrating internal bodily states with neural memory processing. Understanding how visceral oscillators interact with cortical sleep dynamics may therefore reveal new mechanisms linking physiological regulation to objective and subjective sleep experience.

We recorded high-density EEG simultaneously with electrogastrography (EGG) across overnight sleep in 60 healthy adults over two nights (44 female; mean age of 23.1 ± 2.8 years; mean BMI 22.5 ± 3.9 kg/m^2^; Fig. 1A, B). To better understand gastric electrophysiology in healthy human sleep, we assessed three features of the EGG signal: 1) power in the normogastric frequency band (0.033 to 0.067 Hz) (*12*), 2) phase (i.e., the timing within the gastric slow-wave cycle), and 3) amplitude changes of the gastric slow wave relative to canonical sleep events. We observed that these gastric features 1) change dynamically across sleep stages and with the occurrence of key NREM cortical events, and 2) relate to next-day behavioral outcomes, including memory recall and subjective sleep quality.

**Fig. 1.**
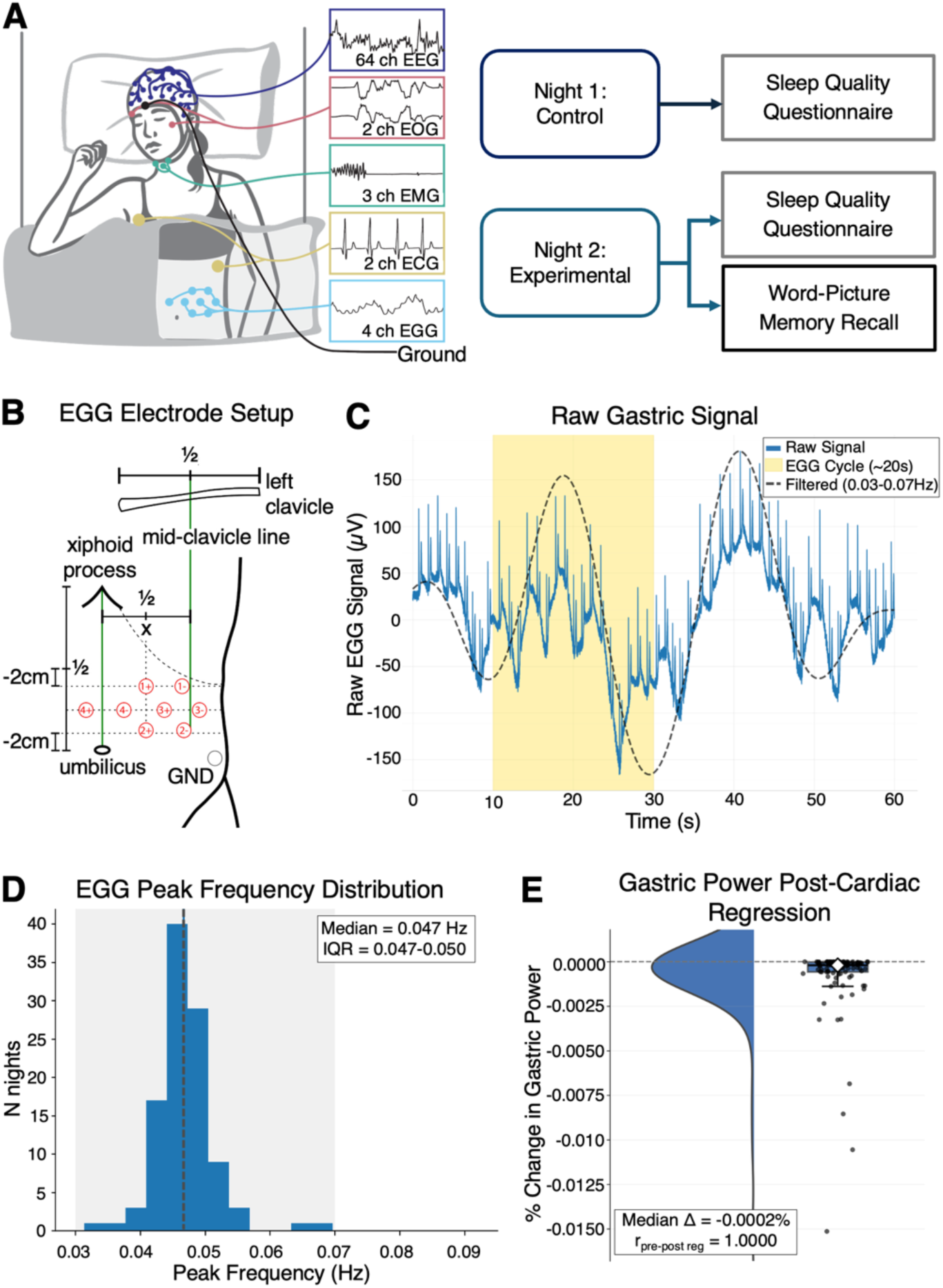
Multimodal sleep recordings capture stable normogastric rhythms during overnight human sleep. **(A)** Simultaneous non-invasive electrophysiological recordings included 64-channel EEG, 2-channel EOG, 3-channel EMG, 2-channel ECG, and 4-channel EGG. Participants reported subjective sleep quality scores and performed a memory recall test on the experimental night. **(B)** Localization of EGG electrode setup for a bipolar montage with respect to anatomical landmarks. Adapted from Wolpert et al. (2020) (CC BY-NC license). **(C)** Representative raw EGG trace. The ∼20 second gastric slow-wave cycle is highlighted (yellow shaded region), with the dashed line indicating the band-pass-filtered gastric rhythm (0.03 to 0.07 Hz). Sharp deflections in the raw signal correspond to cardiac artifacts. **(D)** Group-level distribution of the peak frequency in the overnight EGG signal presents a median value of 0.047 Hz, which lies in the normogastric frequency range (0.03 to 0.07 Hz; n = 105 nights). **(E)** Gastric power remains minimally unchanged (median Δ= −0.0002%) after ECG regression was performed on the EGG signal (n = 105 nights).

## Gastric power dynamically tracks sleep architecture

Preprocessed EGG recordings exhibited a stable gastric slow wave with a period of approximately 20 seconds (Fig. 1C). Across all nights, the peak frequency of this rhythm was tightly distributed around 0.05 Hz (Fig. 1D, and Fig. S1A). This signal primarily reflects gastric electrophysiology: the gastric frequency band accounted for a larger proportion of low-frequency power (<2 Hz) than cardiac-related frequencies (Fig. S1B), showed minimal association with simultaneously recorded ECG signals (Fig. 1E), and exhibited no significant phase-locking with cardiac activity (Fig. S1C). Together, these results confirm that the recorded EGG signal is dominated by gastric activity.

As human sleep unfolds in recurring cycles of NREM and REM sleep across the night, we aimed to understand how gastric electrophysiology varies over the course of the night. Though peak gastric frequency (∼0.05 Hz) was stable across the night (Fig. 2A), gastric band power varied systematically with sleep architecture. Gastric power was significantly greater during NREM than REM sleep (Fig. 2C). Additionally, it was highest early in the night and declined across successive sleep cycles (Fig. 2B and Fig. S2A). This effect was not explained by time since the last reported meal and was consistent across experimental conditions (Fig. S2B, Table 1), with similar trends across sleep stages (Fig. S2C) and sex (Fig. S2D). Together, these findings suggest that gastric power tracks the dynamics of sleep macroarchitecture.

**Fig. 2.**
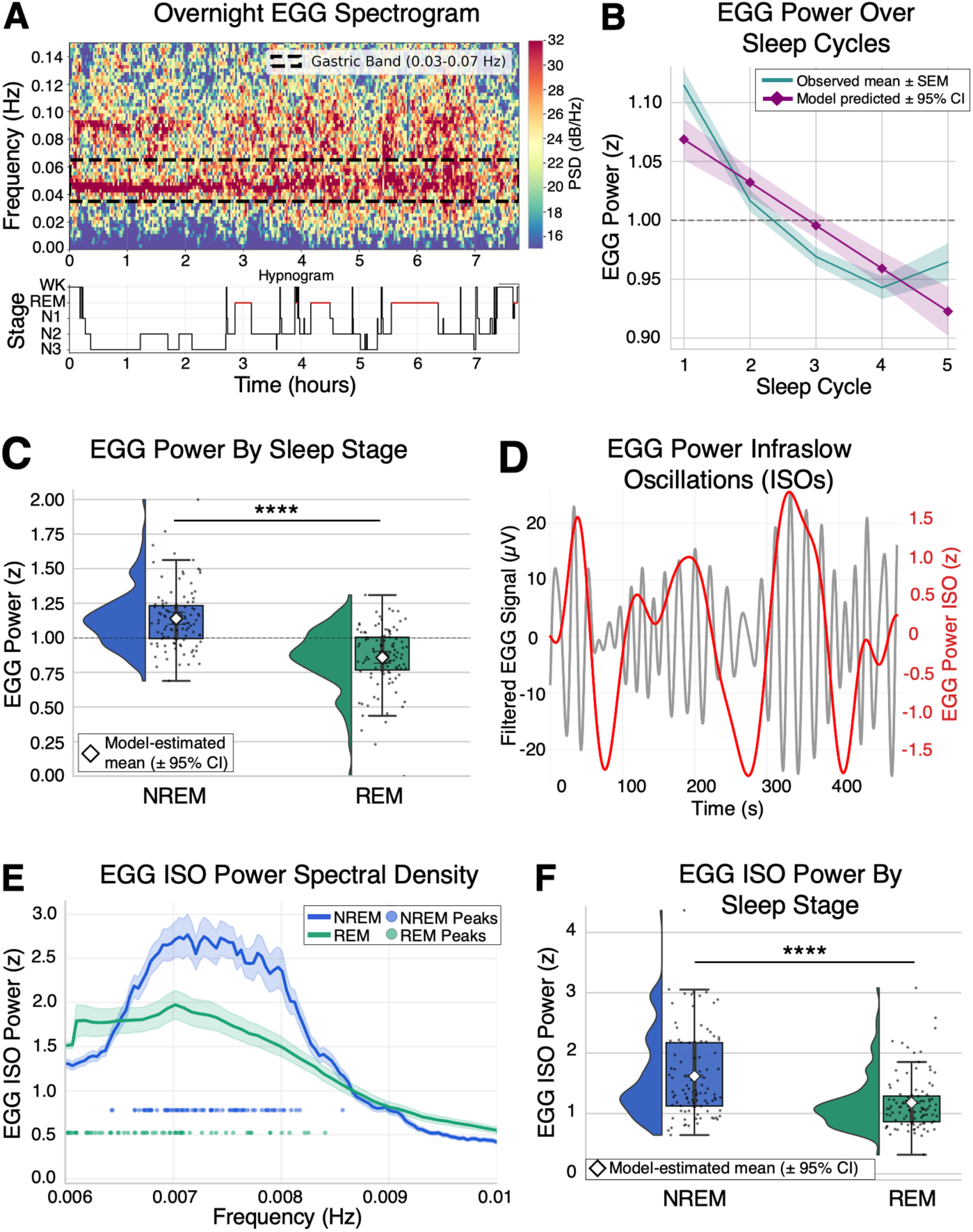
Gastric activity exhibits sleep-stage dependence and infraslow oscillatory structure during NREM sleep. **(A)** Overnight EGG spectrogram in one participant shows sustained power in the normogastric frequency range (dashed box), with the corresponding hypnogram below. **(B)** Gastric power declines significantly across sleep cycles (n = 105 nights). Mean ± SEM (teal) and model-estimated marginal means (EMMs) ± 95% CI (magenta). **(C)** Gastric power was significantly higher during NREM than REM sleep (n = 104 nights, p < 0.0001). Diamonds and error bars indicate EMMs ± 95% CI. **(D)** Example of filtered gastric signal (gray) and corresponding infraslow oscillations (ISOs) in EGG power (red), showing rhythmic fluctuations in gastric amplitude over time. **(E)** EGG ISO power spectra across sleep stages presents a clear infraslow peak (∼0.0075 Hz; period of ∼2 minutes; n = 103 nights), which was more pronounced during NREM than REM sleep. Shaded regions denote ± SEM. **(F)** EGG ISO power across nights with EMMs ± 95 % CI indicate significantly higher ISO power during NREM than REM sleep (n = 97 nights). ns (not significant) p-value ≥ 0.05, \**p*-value *<* 0.05, \*\**p*-value *<* 0.01, \*\*\**p*-value *<* 0.001, \*\*\*\**p*-value *<* 0.0001. All values corrected for multiple comparisons when necessary. See Tables 1-3 for information on statistical analyses and sample sizes.

**Table 1.**
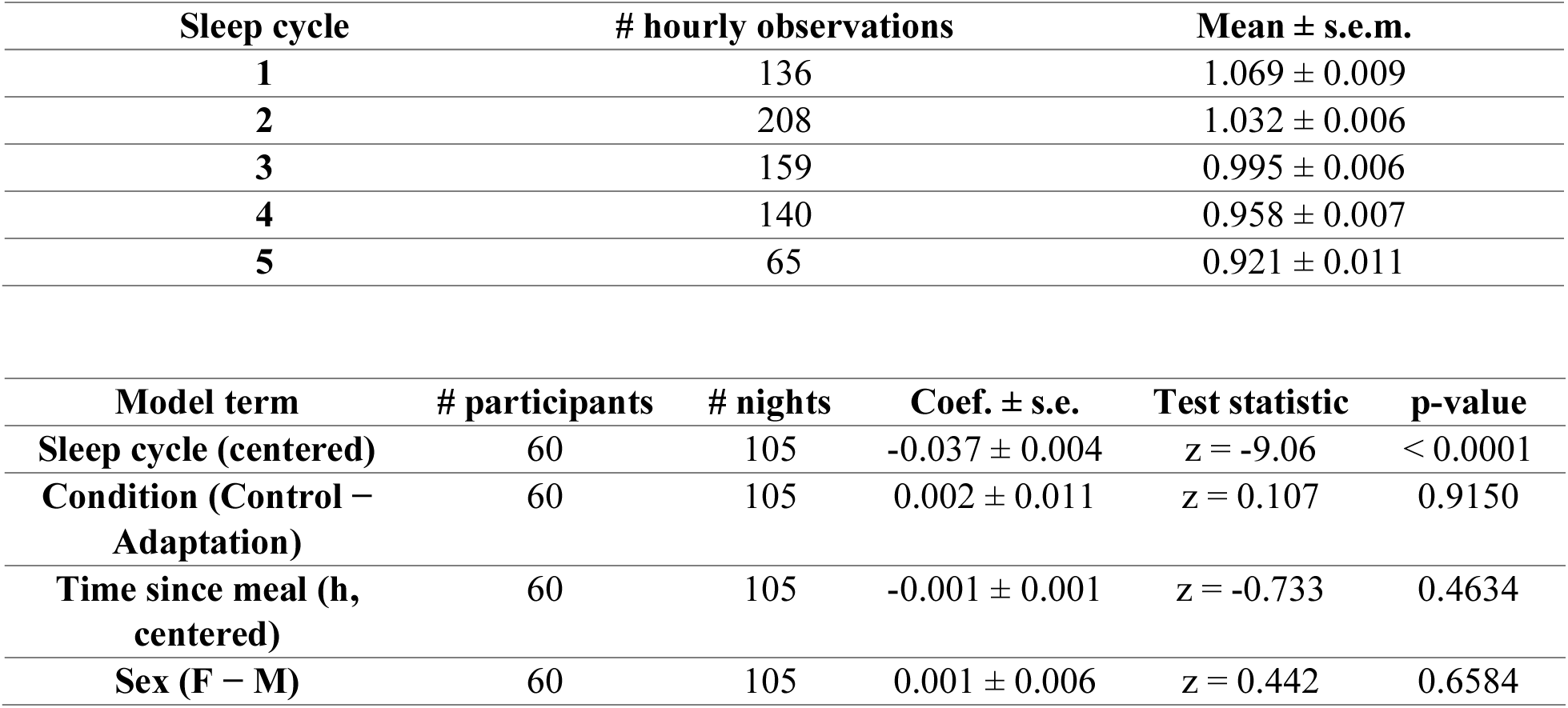
Hourly gastric power across sleep cycles. Displayed values are observed mean ± s.e.m. for cycles 1-5 used in Fig. 2B, Fig. S2A-B. The primary statistical test is the centered sleep-cycle term from a linear mixed-effects model with condition, time since meal, and sex as covariates.

| Sleep cycle | # hourly observations | Mean ± s.e.m. |
| --- | --- | --- |
| 1 | 136 | 1.069 ± 0.009 |
| 2 | 208 | 1.032 ± 0.006 |
| 3 | 159 | 0.995 ± 0.006 |
| 4 | 140 | 0.958 ± 0.007 |
| 5 | 65 | 0.921 ± 0.011 |

| Model term | # participants | # nights | Coef. ± s.e. | Test statistic | p-value |
| --- | --- | --- | --- | --- | --- |
| Sleep cycle (centered) | 60 | 105 | -0.037 ± 0.004 | $z = -9.06$ | $< 0.0001$ |
| Condition (Control – Adaptation) | 60 | 105 | 0.002 ± 0.011 | $z = 0.107$ | 0.9150 |
| Time since meal (h, centered) | 60 | 105 | -0.001 ± 0.001 | $z = -0.733$ | 0.4634 |
| Sex (F – M) | 60 | 105 | 0.001 ± 0.006 | $z = 0.442$ | 0.6584 |

**Table 2.** Normalized gastric power in NREM versus REM sleep. Values are mean ± s.e.m. across nights after within-night normalization from Fig. 2C. The primary statistical test is the stage term from a linear mixed-effects model with condition, time since meal, and sex as covariates.

| Stage | # nights | Mean ± s.e.m. | Statistical test | Test statistic | p-value | Effect size |
| --- | --- | --- | --- | --- | --- | --- |
| NREM | 103 | 1.141 ± 0.022 | Linear mixed-effects model | z = 9.12 | < 0.0001 | d<sub>LMM</sub> = 1.27 |
| REM | 103 | 0.859 ± 0.022 | | | | |
| Comparison | Estimated difference |  | 95% CI | Statistical test |  | p-value |
| NREM vs REM | 0.282 | | [0.221, 0.342] | LMM likelihood-ratio test, $\chi^2(1) = 71.40$ | | < 0.0001 |

**Table 3.** NREM versus REM EGG infraslow oscillation (ISO) peak power. Estimates are from a covariate adjusted linear mixed model fit presented in Fig. 2F. Adjusted means are estimated marginal means with 95% confidence intervals. Covariates included condition, sex, and time since meal stage hour.

| Stage | # nights | Adjusted mean [95% CI] | Statistical test |
| --- | --- | --- | --- |
| NREM | 97 | 1.5772 [1.5633, 1.5917] | Linear mixed model |
| REM | 97 | 1.3505 [1.3366, 1.3651] | Linear mixed model |

| Comparison | Effect size ($\beta$) | Test statistic (z) | Corrected p-value | $d_{LMM}$ |
| --- | --- | --- | --- | --- |
| NREM vs REM | 0.4436 | 5.696 | 0.0000 | 0.817 |

### Gastric power contains an infraslow oscillation amplified during NREM sleep

Given that gastric power varied across sleep stages, we next asked whether it exhibits slower temporal fluctuations. We observed that gastric power exhibited infraslow oscillations (ISOs; Fig. 2D), with a distinct peak near 0.0075 Hz, corresponding to cycles of approximately 2 minutes (Fig. 2E). The strength of this gastric ISO activity was significantly higher during NREM than REM sleep (Fig. 2F and Fig. S3B) and wakefulness (Fig. 2A, B). These observations indicate that gastric electrophysiology during sleep is hierarchically organized across two timescales, in which the 0.05 Hz gastric slow wave is itself modulated by an infraslow rhythm at 0.0075 Hz, both of which are more pronounced during NREM sleep.

### Stomach-brain phase-amplitude coupling varies across NREM sleep stages

Given that gastric-band (∼0.05 Hz) and gastric ISO (∼0.0075 Hz) power were more pronounced during NREM sleep, we asked whether gastric electrophysiology is coordinated with key NREM cortical rhythms, namely slow-wave activity in the delta-band range (0.5 to 4 Hz) and sigma-band activity (10 to 16 Hz) associated with sleep spindles (*6*, *23*). Because these rhythms are differentially expressed across NREM stages, we examined this coordination across N1, N2, and N3 sleep.

Phase-amplitude coupling (PAC) (*24*) allows one to quantify how the phase of a slower signal, in this case the gastric slow wave, is coordinated with the amplitude of a higher frequency signal, or EEG oscillations (Fig. 3A). PAC analyses revealed significant coupling between gastric phase and EEG delta-band amplitude across all 64 EEG channels, as well as between gastric phase and sigma-band amplitude in left frontal and centroparietal regions during N2 sleep (Fig. 3B and Fig. S4). In contrast, these measures were markedly reduced and did not reach significance in N3 (Fig. 3B and Fig. S4). These results suggest that stomach-brain PAC is stage-dependent, with coupling being significant and spatially organized during N2 sleep and comparatively weaker during N3.

**Fig. 3.**
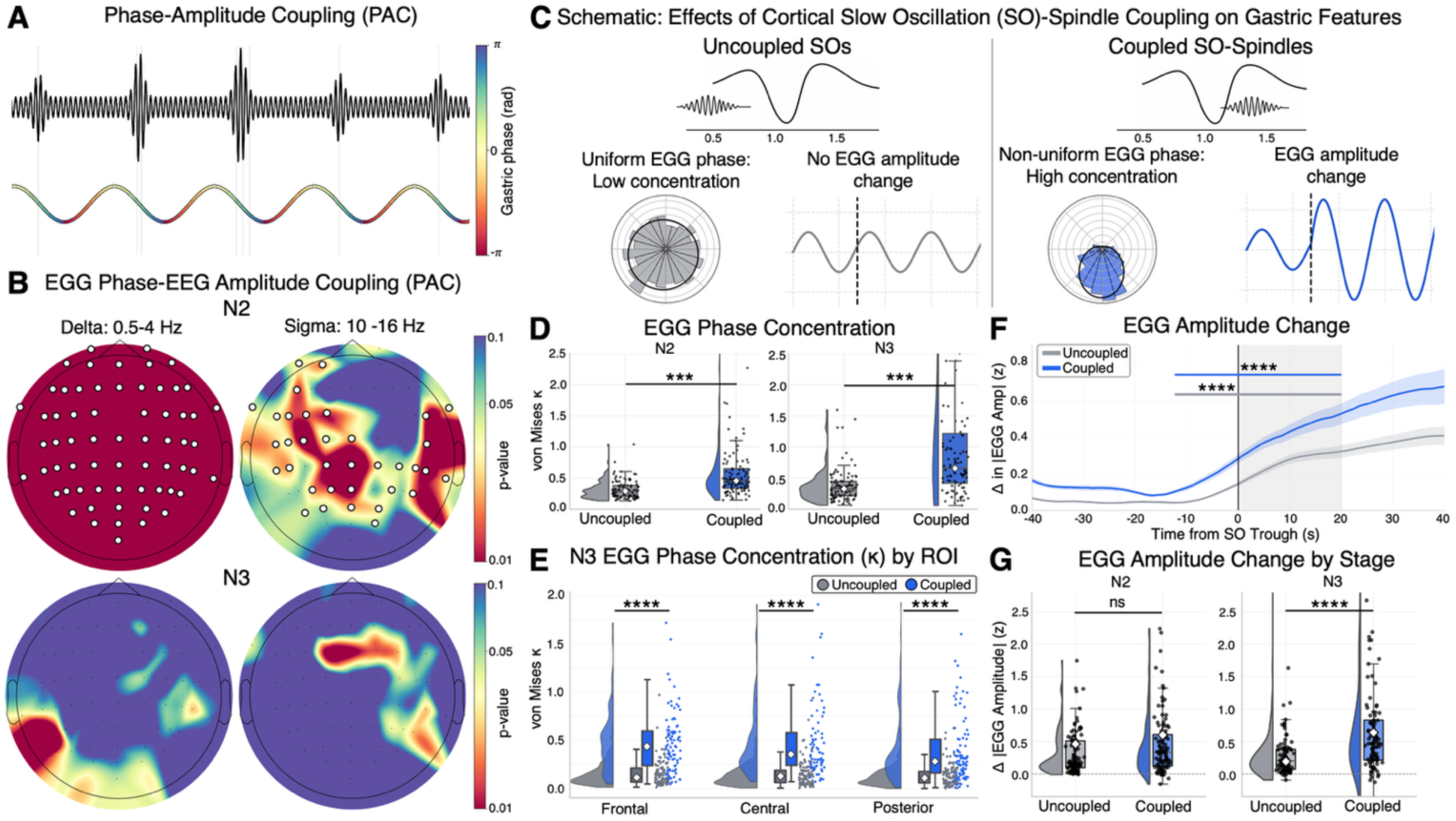
Cortical slow oscillation-spindle events organize gastric phase and amplify gastric activity during sleep. **(A)** Phase amplitude coupling schematic. **(B)** Topographical distribution of EGG phase and EEG delta (left) and sigma (right) amplitude coupling during N2 (top) and N3 (bottom). EGG-EEG phase amplitude coupling is significant and spatially organized in N2 sleep, with global coupling in EEG delta and centro-parietal coupling in EEG sigma (white regions indicate corrected *p <* 0.05; n = 105 nights). **(C)** Schematic of changes in gastric phase and amplitude during coupled SO-spindle and uncoupled SO events. **(D)** Gastric phase concentration (κ) is significantly increased during coupled SO-spindle events than uncoupled ones, with stronger effects in N3 (4-fold increase) than N2 (2-fold increase) sleep (n = 104 nights). **(E)** Regional analysis of N3 SOs show significant increases in gastric phase concentration across frontal, central, and posterior electrodes (n = 105 nights). **(F)** Time course of absolute EGG amplitude change, relative to baseline, aligned to SO troughs. Both coupled and uncoupled SO events result in increased EGG amplitude change, with a significant post-trough increase (n = 105 nights). Shaded regions denote sem. (**G**) EMM-adjusted post-pre SO trough EGG amplitude change is significantly higher during coupled SO events in N3 than N2 sleep (n = 105 nights). ns (not significant) p-value ≥ 0.05, *p-value *<* 0.05, ** p-value *<* 0.01, *** p-value *<* 0.001, **** p-value *<* 0.0001. All values corrected for multiple comparisons when necessary. See Tables 4-7 for information on statistical analyses and sample sizes.

**Table 4.**
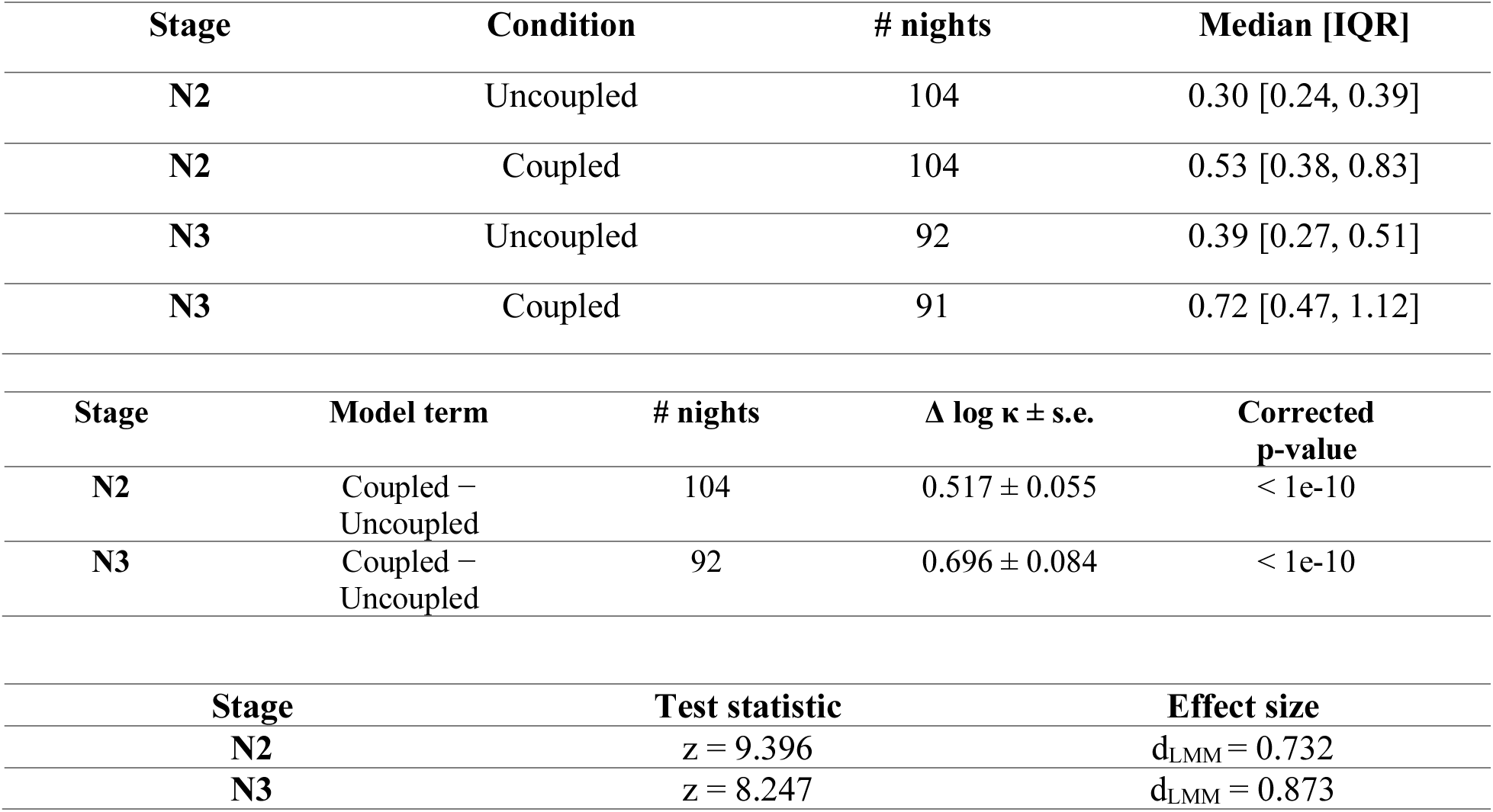
Coupled versus uncoupled EGG phase concentration in N2 and N3. EGG phase concentration (*κ*) during SO events, comparing coupled and uncoupled events separately for N2 and N3. Displayed values in Fig. 3D are per-night median [IQR] of raw *κ*. Statistical tests are covariate adjusted linear mixed-effects models fit on log(*κ*) with sleep cycle, condition, time since meal, and sex as covariates.

### Gastric amplitude and phase are affected by NREM cortical sleep events

Upon observing coupling of the gastric phase to EEG delta and sigma amplitudes in NREM sleep, we sought to understand how gastric electrophysiology changes in relation to discrete cortical events key to NREM. Slow oscillations (SOs) (*<*1 Hz) and spindles (10-16 Hz) are hallmark cortical events in N2 and N3 sleep, reflecting thalamocortical network dynamics that promote neural synaptic plasticity and memory consolidation (*6*, *25*, *26*). While these brain rhythms have been traditionally studied within the context of their roles in intra-and thalamo-cortical processing—such as sensory gating (*2*), coordinating cortical excitability (*27*, *28*), and structuring offline information transfer (*26*, *29*)—the temporal coupling of SOs and spindles coincides with dynamic effects on peripheral physiology such as respiration (*30*, *31*) and cardiac signals (*8*, *32*, *33*). As such, we hypothesized that the occurrence of SO-spindle coupling, defined as a spindle occurring sequentially after the downward phase of the SO (*26*), would be associated with changes in gastric phase and amplitude (Fig. 3C).

To test whether the gastric phase was aligned to the temporal coupling of SOs and spindles, we quantified the strength of phase concentration during the timing of coupled SO-spindles and uncoupled SOs in the EEG signal. Gastric phase concentration was significantly stronger during coupled SO-spindle events than during uncoupled SOs, with increased effects in N3 compared to N2 sleep (Fig. 3D). This coupling effect on gastric phase concentration was sustained through individual sleep cycles (Fig. S5A), and across frontal, central, and posterior EEG regions in both N2 (Fig. S5B) and N3 (Fig. 3E) sleep. Further temporal analysis demonstrated that, when SO-spindle coupling occurred, the gastric phase was more tightly concentrated around the spindle peak than around the SO trough in posterior regions during N2 and across the cortex during N3 (Fig. S5C) sleep. Together, these findings show that the gastric phase is aligned to the timing of coordinated cortical events, with the strongest and most widespread organization during the coupling of spindles and SOs in N3 sleep.

Next, we performed similar analyses to determine whether changes in EGG amplitude were associated with coupled SO-spindle events. We quantified absolute EGG amplitude changes around the trough of coupled SO-spindle events and uncoupled SOs across NREM sleep, and observed that both conditions resulted in significant increases, with a stronger effect during coupled SOs (Fig. 3F). Stage-specific analyses confirmed that gastric amplitude increased significantly after SO troughs in both N2 and N3 (Fig. S6A). However, when directly comparing coupled versus uncoupled SOs within each stage, this increase was significantly greater only in N3, with no significant difference observed in N2 (Fig. 3G, Fig. S6B). Further analysis demonstrated significant gastric amplitude changes aligned with coupled SOs in central EEG regions during N2 and in frontal and posterior regions during N3 (Fig. S6C). Additionally, when SO-spindle coupling occurred, gastric amplitude increases occurred around spindle peaks more than SO troughs in frontal regions during N2 and across all regions in N3 (Fig. 6D).

Altogether, these results suggest that SOs and spindles, traditionally known to facilitate key cortical processes for learning and neuronal plasticity, are also accompanied by coordinated changes in gastric electrophysiology in NREM sleep, revealing regionally specific and event-dependent coupling between cortical dynamics and gastric physiology.

### Gastric power is coupled to infraslow sigma dynamics across NREM sleep

Given that gastric amplitude and phase changes were amplified when spindles accompanied SOs, we next sought to characterize the link between spindle dynamics and gastric activity. Sleep spindles, which lie within the sigma frequency band, occur in 50-second bursts (∼0.02 Hz, Fig.4A) (*7*, *34*). This infraslow oscillation (ISO) of sigma power is a marker of fragility and vigilance during NREM sleep, when our brain, body, and autonomic nervous system (ANS) scan our external environment for potential dangers during sleep (*7*, *8*). Given that the stomach is also influenced by the ANS, we were keen to see whether gastric electrophysiology is coupled to EEG sigma ISOs during NREM sleep.

**Fig. 4.**
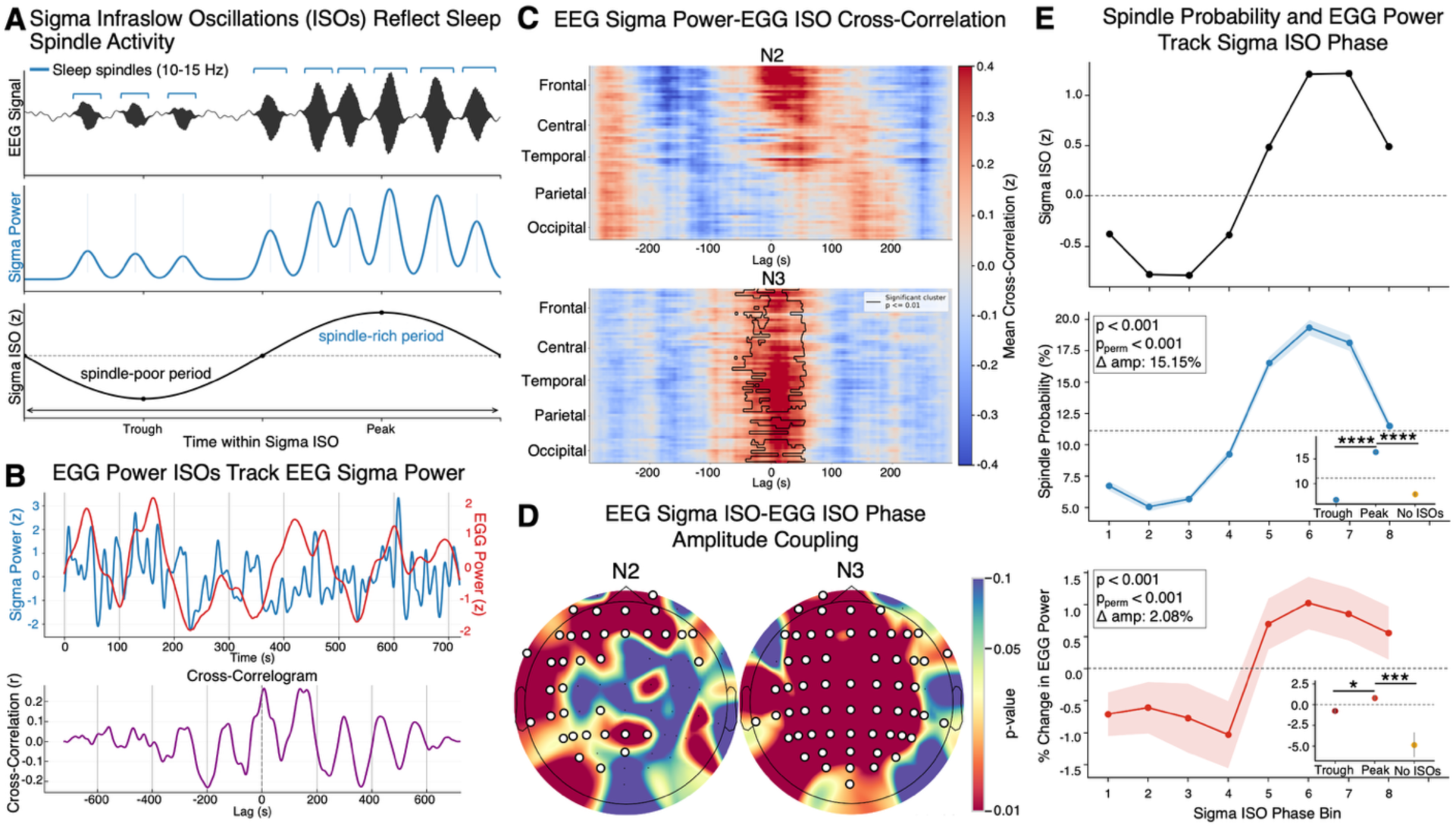
Gastric infraslow dynamics are temporally coupled to cortical sigma oscillations and track spindle activity. **(A)** Schematic describing relationship between sleep spindle activity and EEG sigma ISOs. **(B)** Example trace presenting coordinated infraslow oscillations (ISOs) between EEG sigma power (blue) and EGG gastric power (red) during NREM sleep, with corresponding cross-correlogram below. **(C)** EEG-EGG cross-correlation matrices across all 64 EEG channels during N2 (top) and N3 (bottom) sleep, centered at zero lag for visualization. Significant clusters (*p <* 0.01, cluster-corrected, n = 105 nights) indicate sustained EEG sigma-EGG ISO correlation during N3 sleep. **(D)** Topographical distribution of EGG ISO phase and EEG sigma ISO amplitude coupling during N2 (left) and N3 (right) (highlighted white regions: *p <* 0.05, corrected; n = 105 nights). **(E)** Distribution of sleep spindle probability (middle) and EGG power change (bottom) across phase bins of the infraslow fluctuation of EEG sigma power (top) during NREM sleep. Trough values summarize bins 1-4 and peak values summarize bins 5-8. Both spindle probability and EGG power vary systematically across the sigma ISO cycle, with maximal values near the peak phase. Phase modulation was assessed using sinusoidal (sin(ϕ), cos(ϕ)) model fits and permutation testing (p and p_perm_, respectively). Insets show planned paired contrasts between trough, peak, and no-ISO periods (n = 103 nights). ns (not significant) p-value ≥ 0.05, *p-value *<* 0.05, ** p-value *<* 0.01, *** p-value *<* 0.001, ****p-value *<* 0.0001. All values corrected for multiple comparisons when necessary. See Tables 8 for information on statistical analyses and sample sizes.

Characterization of the sigma ISO confirmed that EEG sigma-band activity is present in both N2 and N3 sleep but is significantly reduced in N3 relative to N2 (Fig. S7B). Despite this, the relative prominence of the ∼0.02 Hz sigma ISO peak was comparable in both N2 and N3 (Fig. S7C), indicating that sigma ISO dynamics persist in deep sleep. These findings are consistent with prior work demonstrating infraslow (∼0.01 to 0.02 Hz) modulation of sigma and spindle activity across NREM sleep, albeit with reduced magnitude in deeper stages (*8*, *34*).

To better match the slow timescales of sigma ISOs, we focused on changes in gastric power rather than transient gastric phase and amplitude changes, which we previously characterized as having a peak frequency of ∼0.0075 Hz, with the strongest signal in NREM sleep (Fig. 2E, F). We observed that EGG power ISOs track EEG sigma power in cross-correlation analyses (Fig. 4B). When extending this analysis to all 64 EEG channels and sleep stages, we observed strong gastric power coupling with infraslow frontal EEG regions during N2 and significant correlation during N3 in all EEG regions (Fig. 4C). Similar analyses in N1, REM, and wake bouts revealed no significant temporal structure (Fig. S8A, B). These results indicate that gastric power is temporally coupled to infraslow sigma dynamics specifically during NREM sleep, with coordinated fluctuations emerging in N2 and becoming globally organized in N3.

While cross-correlation between two signals tests for lagged co-fluctuations, PAC analysis allows us to specifically assess whether the phase of the EGG power is coordinated with the amplitude of EEG sigma power (sigma ISO), over the gastric cycle. PAC analysis between sigma ISO and EGG ISO phase demonstrated significant coupling in frontal, left temporal, and parietal regions during N2, and all cortical regions during N3 (Fig. 4D and Fig. S9). These results demonstrate that gastric power fluctuations are not only temporally correlated with sigma ISOs but are coupled to sigma power in a phase-dependent manner, indicating coordinated brain-body infraslow dynamics across NREM sleep.

### Gastric power is phase-locked to the infraslow sigma cycle and spindle dynamics

Because sigma ISOs and sleep spindles index infraslow fluctuations in NREM fragility and stability, we next asked whether gastric dynamics are temporally aligned to specific phases of this cycle. To this end, we examined how gastric power is organized within the infraslow sigma cycle and across spindle occurrences. We split the sigma ISO cycle (Fig. 4A) into eight equal phase bins, in which we calculated the probability of sleep spindle events and the change in EGG power. By definition, the probability of spindle occurrence was highest during the rise of the sigma ISO cycle (Fig. 4E) (*7*). Similarly, the greatest percent change in gastric power occurred during the same phase, aligning with peaks in spindle probability and sigma power (Fig. 4E). These changes were significant when compared to the trough of the sigma ISO cycle, or spindle-poor periods and bouts of no sigma ISOs, indicating that these changes were aligned with the rise of the sigma ISO cycle (Fig. 4E).

These patterns were consistent for spindle probability across both N2 (Fig. S10A) and N3 (Fig. S10B) sleep. For gastric power, a similar phase-dependent pattern was observed across stages, although statistical significance was primarily driven by N3 sleep, where larger gastric power changes were observed (Fig. S10B). These findings show for the first time that gastric power is temporally aligned with infraslow cycles of sigma power and spindle occurrence in the brain.

### Overnight gastric dynamics relate to memory consolidation and subjective sleep quality

Our discovery of robust stomach-brain coupling during NREM sleep prompted us to ask whether this coordination is associated with functional outcomes relevant to sleep, namely sleep-dependent memory consolidation and sleep quality.

On the experimental night, participants were asked to learn a word-picture association task prior to sleep and recall it the following morning (*35*). The task consisted of 99 word-picture associations pairing neutral words with positive, neutral, and negative pictures, and yielding a percentage-correct recall score. We calculated residual memory recall scores based on the difference between an individual’s baseline performance during the evening recall and the amount recalled the following morning.

We first performed an exploratory screen testing gastric-based features—including sigma-band PAC, sigma power ISO PAC, and gastric phase concentration during N2 and N3 cortical events— against memory and sleep outcomes after residualizing covariates (Fig. S11A, C). Several nominal associations were observed, particularly involving sleep-stage differences in EGG-EEG sigma coupling and N3 gastric phase concentration, which motivated focused confirmatory analyses of memory consolidation and subjective sleep quality.

We first examined whether the relative balance of stomach-brain coupling across NREM stages was related to overnight memory consolidation, motivated by evidence that N2 and N3, which are dominated by spindles and slow oscillations, respectively, make complementary contributions to sleep-dependent memory processing. Stronger EEG sigma-EGG PAC during N2 was associated with better next-morning memory recall (8.9% of variance explained), whereas relatively stronger coupling during N3 than N2 (14.4% of variance explained) was associated with poorer recall after adjusting for SO-spindle coupling strength and spindle density (Fig. 5A). These findings indicate that the functional significance of stomach-brain coupling differs across NREM stages, with relatively stronger sigma-EGG coupling during deeper N3 sleep associated with worse memory outcomes, whereas coupling during lighter N2 sleep is associated with better outcomes.

**Fig. 5.**
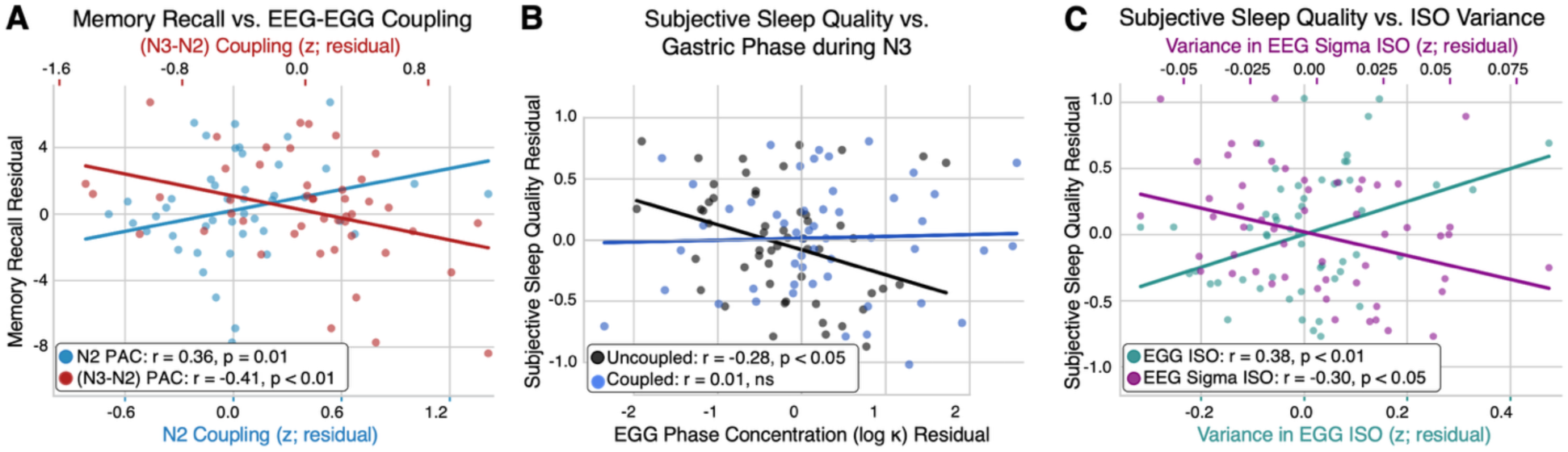
Stomach-brain coupling across timescales differentially predicts memory consolidation and subjective sleep quality. **(A)** Stronger N2 EEG sigma–EGG PAC (blue; r = 0.36, p = 0.01) was significantly associated with improved next-day memory recall, whereas stronger N3–N2 EEG sigma-EGG PAC (red; r = - 0.41, p < 0.01) was significantly associated with worse memory recall (n = 47 nights). Memory recall scores were residualized on EEG-derived measures associated with memory consolidation, including SO-spindle coupling strength and spindle density. **(B)** Greater gastric phase concentration during uncoupled SOs in N3 (black; r = - 0.28, p < 0.05) was associated with lower subjective sleep quality, whereas no significant association was observed for coupled SOs (blue; r = 0.01, p = ns; n = 56 nights). **(C)** Variance in EGG ISO was significantly positively associated with subjective sleep quality (teal; r = 0.38, p < 0.01), whereas variance in EEG sigma ISO was significantly positively associated with subjective sleep quality (magenta; r =-0.30, p < 0.05; n = 56 nights). Subjective sleep quality scores were residualized on EEG-derived measures of sleep architecture commonly used as objective markers of sleep quality. All residualizations were performed using Huber robust regression, and partial Spearman correlations were used for visualization and summary statistics. See Tables 9, 11–13 for statistical details and sample sizes.

**Table 5.** ROI comparison of coupled versus uncoupled SO events in N3. EGG phase concentration (*κ*) by ROI during SO events in N3, comparing coupled and uncoupled events. Displayed values in Fig. 3E are per-night median [IQR] of raw *κ*. Statistical tests are simple effects from a covariate-adjusted linear mixed-effects model fit on log(*κ*) with ROI, sleep stage, coupling, condition, sex, and time since meal as predictors.

| ROI | (# nights) Median<br>uncoupled [95% CI] | (# nights) Median<br>coupled [95% CI] | $\beta \pm \text{s.e.}$ | Test stat.<br>(z) | Corrected p-value |
| --- | --- | --- | --- | --- | --- |
| **Frontal** | (104) 0.13<br>[0.08, 0.22] | (93) 0.41<br>[0.23, 0.66] | $1.133 \pm 0.111$ | 10.242 | $< 1\text{e-}10$ |
| **Central** | (102) 0.15<br>[0.10, 0.28] | (65) 0.45<br>[0.30, 0.71] | $1.137 \pm 0.123$ | 9.248 | $< 1\text{e-}10$ |
| **Posterior** | (104) 0.13<br>[0.08, 0.23] | (92) 0.36<br>[0.18, 0.63] | $1.220 \pm 0.109$ | 11.229 | $< 1\text{e-}10$ |

| ROI | Fold change |
| --- | --- |
| **Frontal** | $\times 3.10$ |
| **Central** | $\times 3.12$ |
| **Posterior** | $\times 3.39$ |

**Table 6.** NREM event-triggered EGG amplitude change. Displayed values in Fig. 3F are raw night-level mean ± s.e.m. of baseline-referenced absolute EGG amplitude in the displayed pre (−40 to 0 s) and post (0 to 20 s) windows during pooled NREM SO events. Statistical tests are within-coupling paired t-tests comparing post versus pre windows.

| **Coupling** | **# nights** | **Pre mean**<br>**± s.e.m.** | **Post mean**<br>**± s.e.m.** | **Estimated**<br>**difference** | **Test stat.**<br>**(z)** | **p-value** | **Effect size** |
| --- | --- | --- | --- | --- | --- | --- | --- |
| **Uncoupled** | 105 | 0.029 ±<br>0.002 | 0.273 ±<br>0.023 | 0.244 | 7.233 | < 0.0001 | $d_{LMM} =$<br>0.998 |
| **Coupled** | 105 | 0.076 ±<br>0.006 | 0.482 ±<br>0.045 | 0.406 | 12.019 | < 0.0001 | $d_{LMM} =$<br>1.659 |

**Table 7.** Stage-specific post-pre change in event-triggered EGG amplitude. Stage-specific observed post-pre change in baseline-referenced absolute EGG amplitude for coupled and uncoupled SO events. Displayed values in Fig. 3G are raw night-level mean ± s.e.m. by stage and coupling condition. The primary statistical test was the stage-specific coupled-uncoupled contrast from a linear mixed-effects model with event count, experimental condition, sex, time since last meal, and normalized delta power as covariates, with repeated measures accounted for by subject and night random effects.

| Stage | Uncoupled mean<br>± s.e.m. | Coupled mean<br>± s.e.m. | Estimated<br>difference | Test stat. | p-value | Effect size |
| --- | --- | --- | --- | --- | --- | --- |
| **N2** | 0.316 ± 0.032 | 0.453 ± 0.046 | 0.137 | $z = 1.605$ | 0.1084 | $d_{LMM} = 0.223$ |
| **N3** | 0.352 ± 0.066 | 0.799 ± 0.127 | 0.447 | $z = 5.189$ | < 0.0001 | $d_{LMM} = 0.727$ |

**Table 8.** Shared sigma ISO phase modulation during pooled NREM sleep. Shared sigma ISO phase modulation of spindle probability and EGG power during NREM from Fig. 4E. Trough values summarize bins 1-4, peak values summarize bins 5-8, and no-ISO values summarize samples outside detected sigma ISOs. Omnibus phase-modulation *p*-values are from mixed-effects models with sin(φ) and cos(φ) terms fit across bins 1–8. Permutation p-values are based on 1000 within-unit phase shuffles.

| Outcome | # units | Trough mean ± s.e.m. | Peak mean ± s.e.m. | No ISOs mean ± s.e.m. |
| --- | --- | --- | --- | --- |
| Spindle probability (%) | 97 | $6.68 \pm 0.22$ | $16.37 \pm 0.23$ | $7.82 \pm 0.38$ |
| EGG power change (%) | 103 | $-0.78 \pm 0.32$ | $0.78 \pm 0.32$ | $-4.87 \pm 1.49$ |

| Outcome | Statistical test | Omnibus $p$-value | Permutation $p$-value | Peak-to-trough amp (%) |
| --- | --- | --- | --- | --- |
| Spindle probability (%) | Mixed model ($\sin\phi$, $\cos\phi$) | $5.66 \times 10^{-232}$ | $< 0.001$ | 15.15 |
| EGG power change (%) | Mixed model ($\sin\phi$, $\cos\phi$) | $9.90 \times 10^{-6}$ | $< 0.001$ | 2.08 |

| Outcome | Comparison | Holm-corrected $p$-value |
| --- | --- | --- |
| Spindle probability (%) | Peak vs Trough | $1.68 \times 10^{-38}$ |
| Spindle probability (%) | Peak vs No ISOs | $1.47 \times 10^{-30}$ |
| EGG power change (%) | Peak vs Trough | 0.010 |
| EGG power change (%) | Peak vs No ISOs | $8.96 \times 10^{-4}$ |

**Table 9.**
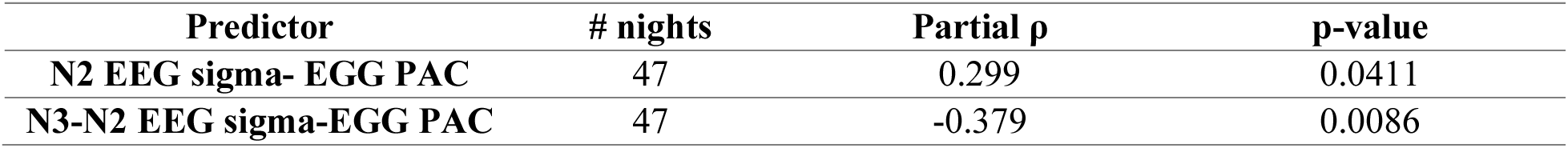
Combined stage-specific phase-amplitude coupling figure statistics. Morning memory was residualized on sex, time since meal, stage-specific N2/N3 CO-spindle coupling, and stage-specific N2/N3 spindle density, then plotted in the same adjusted space against N2 gastric– sigma coupling and stage-dependent gastric-sigma coupling. Displayed statistics in Fig. 5A are partial Spearman correlations from the Huber-robust figure analyses.

| Predictor | # nights | Partial $\rho$ | p-value |
| --- | --- | --- | --- |
| N2 EEG sigma- EGG PAC | 47 | 0.299 | 0.0411 |
| N3-N2 EEG sigma-EGG PAC | 47 | -0.379 | 0.0086 |

**Table 10.** SF-AR sleep questionnaire items used to compute the subjective sleep quality score. Reverse-scored items are indicated in parentheses.

| Question number | Item wording |
| --- | --- |
| 1 | I slept restfully |
| 2 | I slept well |
| 3 | I slept deeply |
| 4 | I woke up feeling refreshed |
| 5 | I woke up feeling rested |
| 6 | I fell asleep easily |
| 7 | I woke up easily |
| 8 | I had uninterrupted sleep |
| 9 | I slept without restlessness |
| 10 | I felt awake |
| 11 | I felt able to concentrate |
| 12 | I felt balanced |
| 13 | I felt fit |
| 14 | I felt carefree |
| 15 | I felt good about my dreams |
| 16 | I did not have disturbing dreams |
| 17 | My dreams were pleasant |
| 18 | I felt dozy (reverse-scored) |
| 19 | I felt exhausted (reverse-scored) |
| 20 | I felt in need of sleep (reverse-scored) |
| 21 | I felt tired (reverse-scored) |
| 22 | I felt unable to cope (reverse-scored) |

**Table 11.**
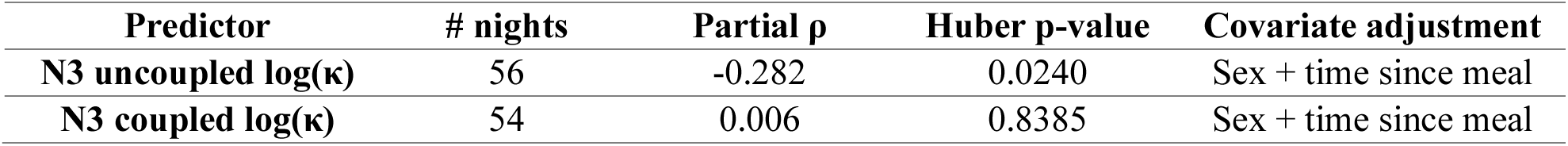
Subjective sleep quality association with N3 coupling strength. Subjective sleep quality in control nights was residualized on sex and time since meal, then related to N3 coupling strength separately for uncoupled and coupled cycles. Reported correlation in Fig. 5B are partial Spearman ρ in the adjusted space, and inferential p-values are from the corresponding Huber-robust regressions.

| Predictor | # nights | Partial $\rho$ | Huber p-value | Covariate adjustment |
| --- | --- | --- | --- | --- |
| N3 uncoupled $\log(\kappa)$ | 56 | -0.282 | 0.0240 | Sex + time since meal |
| N3 coupled $\log(\kappa)$ | 54 | 0.006 | 0.8385 | Sex + time since meal |

**Table 12.** Control-night comparison of gastric and cortical predictors of subjective sleep quality. Subjective sleep quality in control nights was residualized on sleep architecture, sex, and time since meal, then related to intrinsic EGG ISO variability and EEG sigma ISO variability. Reported correlations in Fig. 5C are partial Spearman ρ in the adjusted space, and inferential p-values are from the corresponding Huber-robust regressions. No condition covariate was included because only control nights were analyzed.

| Predictor | # nights | Partial $\rho$ | Huber p-value | Covariate adjustment |
| --- | --- | --- | --- | --- |
| **EGG ISO amplitude SD** | 56 | 0.378 | 0.0194 | PSG + sex + meal time |
| **EEG sigma ISO amplitude SD** | 56 | -0.304 | 0.0197 | PSG + sex + meal time |

**Table 13.** Omnibus subjective sleep quality model with physiological predictors. A base PSG architecture model for subjective sleep quality was compared with a full model adding EGG ISO variability, EEG sigma ISO variability, and N3 uncoupled coupling.

| Model | # nights | Predictors | R<sup>2</sup> | Adjusted R<sup>2</sup> | Stat. test |
| --- | --- | --- | --- | --- | --- |
| PSG-only | 56 | Architecture + sex + meal time | 0.165 | 0.001 | Reference |
| PSG + physiology | 56 | + EGG ISO + EEG sigma ISO + N3 uncoupled κ | 0.369 | 0.193 | F(3,43) = 4.643, p = 0.0067 |

| Physiological term | Standardized coefficient | p-value |
| --- | --- | --- |
| EGG ISO variability | 0.232 | 0.005 |
| EEG sigma ISO variability | -0.155 | 0.039 |
| N3 uncoupled coupling | 0.109 | 0.179 |

Using data from the control night, we next examined whether gastric coupling measures were associated with subjective sleep quality. Regression analyses showed that stronger gastric phase concentration during N3 uncoupled SOs was significantly associated with poorer subjective sleep quality, whereas gastric phase concentration during spindle-coupled SOs showed no significant relationship with subjective sleep quality (Fig. 5B). These results show that gastric phase alignment relative to specific cortical events differs between conditions, with N3 coupling outside of spindle-associated activity associated with poorer subjective sleep quality.

Finally, we asked whether an EGG-only intrinsic measure, namely the infraslow variability of gastric power, is associated with sleep quality. Higher variability in gastric ISO amplitude was positively associated with subjective sleep quality, whereas higher variability in EEG sigma ISO amplitude was negatively associated with subjective sleep quality in the covariate-adjusted model (Fig. 5C). Together, gastric ISO variability, EEG sigma ISO variability, and N3 uncoupled gastric phase concentration increased the explained variance in subjective sleep quality by 20.4% beyond standard polysomnographic and demographic covariates (Table 13), suggesting a substantial incremental contribution of gastric dynamics. This increase was distributed across gastric ISO variability (12.8%), EEG sigma ISO variability (6.3%), and N3 uncoupled gastric phase concentration (1.3%). Our further analysis revealed that gastric ISO variance significantly outperformed its association with subjective sleep quality scores when compared with a commonly used cardiac metric, heart rate variability (*36*), for assessing subjective sleep quality (Fig. S11C).

In contrast to the stomach-brain coupling measures, these findings indicate that intrinsic gastric dynamics, independent of their alignment to cortical events, are positively associated with subjective sleep quality.

Together, our findings indicate that gastric physiology during sleep is unlikely to be epiphenomenal. Rather, the functional relevance of stomach-brain interactions during sleep is context-dependent: stronger or relatively enhanced coupling during deep NREM sleep is associated with poorer memory performance and reduced subjective sleep quality, independent of cortical and cardiac measures.

## Discussion

Sleep has increasingly been recognized as a state of continuous brain-body communication, yet whether and how gastric rhythms participate in human sleep physiology remains almost completely unknown. Here, by combining high-density EEG with overnight EGG, we show that gastric electrophysiology is not merely background noise but changes dynamically throughout the night. Gastric electrophysiology tracks sleep macroarchitecture, remaining sustained throughout the night and elevated during NREM sleep, while exhibiting hierarchical temporal organization in which the canonical ∼0.05 Hz gastric slow wave is modulated by a minute-scale infraslow rhythm that is strongest during NREM. Within this framework, stomach-brain coupling is not uniform but exhibits stage-and event-specificity. Gastric phase is coordinated with cortical delta and sigma activity during N2 sleep but attenuated in deeper N3 sleep, and slow oscillation-spindle complexes are accompanied by systematic changes in gastric phase and amplitude. At slower timescales, gastric power fluctuations are phase-locked to the infraslow sigma cycle, peaking during phases of high spindle probability, indicating that gastric dynamics are embedded within ongoing cortical infraslow organization. While sleep is often characterized as a state in which the brain is functionally disconnected from the external environment (*2*), our findings show that it remains dynamically coupled to internal physiological signals. Furthermore, interoceptive dynamics during sleep may influence memory consolidation and subjective sleep experience, consistent with growing evidence that visceral signals shape cognitive and affective processes during wakefulness (*21*). We show that the functional relevance of these interactions is context-dependent: stomach-brain coupling during lighter N2 sleep is associated with better memory outcomes, whereas relatively stronger coupling during deeper N3 sleep and outside spindle-associated activity is linked to poorer memory and reduced subjective sleep quality. Additionally, stable gastric infraslow dynamics predict better sleep quality independent of cortical and cardiac measures. As such, our findings demonstrate that gastric electrophysiology during sleep is structured, selectively coordinated with cortical dynamics, and behaviorally meaningful. More importantly, they indicate that the association between gastric physiology, cognition, and sleep experience depends critically on when during NREM sleep, and in relation to which cortical events gastric signals are integrated—challenging the prevailing view of sleep as a predominantly brain-centered state.

Prior work on brain-heart (*33*) and brain-respiration (*29*) coupling has shown that central and autonomic rhythms form a jointly organized system during NREM sleep, with stage-dependent changes in coupling strength and links to arousal and homeostasis. By demonstrating that gastric rhythms are similarly structured, stage-dependent, and integrated with canonical cortical oscillations, we extend this framework to include visceral signals from the gut, suggesting that cortical events in NREM sleep serve not only as windows for cortico-thalamic communication but also for cortico-visceral coordination. We show that slow oscillation-spindle complexes, which provide temporally precise “communication windows” for hippocampal-neocortical information transfer (*37*, *38*), are accompanied by systematic changes of gastric phase and amplitude. This suggests that slow oscillations and spindles help structure the timing and magnitude of stomach-brain interactions, potentially influencing both neural replay and the alignment of central processing with internal physiological state (*39*). At slower timescales, our findings further suggest that visceral and thalamo-cortical systems share common organizing dynamics during NREM sleep. The observation that the gastric slow wave is modulated by a slower envelope paralleling infraslow modulation of sigma power and spindle activity on similar timescales (*7*, *8*). Coupling between gastric and sigma infraslow rhythms, therefore, appears to embed the stomach within the alternation of “fragile” and “protected” NREM microstates linked to arousability and noradrenergic activity (*7*, *8*). In this view, our results suggest that gastric-sigma coupling is stronger during these protected periods of low arousability and weaker during fragile periods when the sleeping brain is more susceptible to awakening. Together, these findings suggest that brain-body coupling during sleep is dynamically organized across events, stages, and timescales.

By leveraging continuous electrophysiology, we here uncover fast, event-locked, and infraslow stomach-brain coupling modes and place gastric activity within a neural framework aligned with key sleep oscillations. Several pathways could mediate this coordination. Vagal afferents relay gastric signals to the nucleus tractus solitarius, which projects to locus coeruleus, parabrachial nuclei, thalamus, and insula (*40*), providing routes through which gastric rhythms could influence cortical excitability. Descending autonomic pathways may impose a common infraslow structure on both gastric motility and cortical sigma activity, creating reciprocal loops through which sigma ISOs periodically sample internal bodily state. In parallel, spinal afferents via the splanchnic nerve modulate locus coeruleus activity during gastric distension and, through efferent pathways, alter gastric pacemaker activity (*41*), suggesting that both afferent and efferent noradrenergic pathways contribute. Another possibility is that shared infraslow hemodynamic fluctuations contribute to the observed coupling. Simultaneous EEG-fMRI studies demonstrate that infraslow vascular dynamics are coordinated with NREM sleep slow waves and autonomic arousal indices (*42*, *43*), motivating the hypothesis that gastric and cortical signals are co-embedded within a global arousal-related neurovascular and autonomic physiological rhythm.

Our analyses are inherently correlational, precluding inference about directionality. Addressing directionality will likely require passive causal inference methods, such as Granger causality (*44*) or directed information (*45*), applied in datasets specifically designed to test whether information flow is preferentially stomach-to-brain, brain-to-stomach, or bidirectional across NREM contexts. Active manipulations, such as gastrointestinal stimulation, vagus nerve stimulation (*46*), or cortical perturbation (*47*), could further test causal interactions within this system. Methodologically, surface EGG recordings limit spatial resolution and do not capture propagation dynamics of the gastric slow wave. Future studies using high-resolution EGG (*48*), source localization (*49*), or invasive recordings (*50*) could resolve regional differences in gastric activity. Furthermore, incorporating additional peripheral measures, such as a respiratory belt, would enable testing of how gastric, cardiac, and respiratory rhythms jointly interact with cortical dynamics during sleep. Finally, our framework generates testable predictions about where targeted modulation of gastric activity (e.g., meal timing, pharmacology, or peripheral stimulation) can alter specific coupling modes and produce measurable changes in memory, arousability, and subjective sleep quality. In conclusion, our results argue that sleep is a coordinated, multi-organ physiological state in which central and visceral rhythms are jointly structured, motivating the incorporation of gastric physiology into future sleep studies to better understand the mechanisms underlying sleep restoration and cognitive function.

## Acknowledgments

We thank Nico Adelhöfer, Somayeh Ataei, Lucija Blazevski, Lisa Hazebrouqc, Sicelo Jones, Stergiani Lentzou, Seungwon Nam, Anastasiya Paltarhhitskaya, Famke Roest, Yibran Pacheco Saez, Sofia Tzioridou, Tinke Vanbuijtenen, Floris van der Werff, Zora Wesseley, Selin Yavuzcan, Jialin Zhao for help with the data collection. We thank Mackenzie Fredericks, Catherine Liu and Gladys Ornelas for help with data preprocessing. We thank Stephen Matheson for help with the manuscript.

## Author contributions

SFS designed the experiment and collected data. AAR analyzed data and wrote the manuscript. All authors reviewed and edited the manuscript.

## Data, code, and materials availability

This study comprises the analysis of the dataset from the following pre-registered report (*51*). All processed data and source data supporting the findings of this study are publicly available at the Stanford Digital Repository (*52*). As publicly sharing the raw data is prohibited due to participant confidentiality and ethics protocols, data may be shared upon request. To obtain data, please contact the corresponding author, Todd Coleman, or first author, Akshita Rao.

## Supplementary Materials

Materials and Methods

Figs. S1 to S11

Tables S1 to S23

References (*53–65*)

## References and Notes

1. M. G. Frank, H. C. Heller, “The Function(s) of Sleep” in Sleep-Wake Neurobiology and Pharmacology, H.-P. Landolt, D.-J. Dijk, Eds. (Springer International Publishing, Cham, 2019), pp. 3–34.

2. A. Coenen, Sensory gating and gaining in sleep: the balance between the protection of sleep and the safeness of life (a review). J. Sleep Res. 33, e14152 (2024).

3. M. de Zambotti, J. Trinder, A. Silvani, I. M. Colrain, F. C. Baker, Dynamic coupling between the central and autonomic nervous systems during sleep: A review. Neurosci. Biobehav. Rev. 90, 84–103 (2018).

4. L. K. McCorry, Physiology of the autonomic nervous system. Am. J. Pharm. Educ. 71, 78 (2007).

5. A. M. Fink, U. G. Bronas, M. W. Calik, Autonomic regulation during sleep and wakefulness: a review with implications for defining the pathophysiology of neurological disorders. Clin. Auton. Res. 28, 509–518 (2018).

6. M. Steriade, D. A. McCormick, T. J. Sejnowski, Thalamocortical oscillations in the sleeping and aroused brain. Science 262, 679–685 (1993).

7. A. Osorio-Forero, R. Cardis, G. Vantomme, A. Guillaume-Gentil, G. Katsioudi, C. Devenoges, L. M. J. Fernandez, A. Lüthi, Noradrenergic circuit control of non-REM sleep substates. Curr. Biol. 31, 5009–5023.e7 (2021).

8. S. Lecci, L. M. J. Fernandez, F. D. Weber, R. Cardis, J.-Y. Chatton, J. Born, A. Lüthi, Coordinated infraslow neural and cardiac oscillations mark fragility and offline periods in mammalian sleep. Sci. Adv. 3, e1602026 (2017).

9. L. S. Bueno-Junior, M. S. Ruckstuhl, M. M. Lim, B. O. Watson, The temporal structure of REM sleep shows minute-scale fluctuations across brain and body in mice and humans. Proc. Natl. Acad. Sci. 120, e2213438120 (2023).

10. T. Väyrynen, H. Helakari, V. Korhonen, J. Tuunanen, N. Huotari, J. Piispala, M. Kallio, L. Raitamaa, J. Kananen, M. Järvelä, J. Matias Palva, V. Kiviniemi, Infra-slow fluctuations in cortical potentials and respiration drive fast cortical EEG rhythms in sleeping and waking states. Clin. Neurophysiol. 156, 207–219 (2023).

11. X. Yang, F. T. Fedumenti, N. Niethard, M. Hallschmid, J. Born, K. Rauss, Regulation of peripheral glucose levels during human sleep. *Sleep*, zsaf042 (2025).

12. N. Wolpert, I. Rebollo, C. Tallon-Baudry, Electrogastrography for psychophysiological research: Practical considerations, analysis pipeline, and normative data in a large sample. Psychophysiology 57, e13599 (2020).

13. K. M. Sanders, S. D. Koh, S. M. Ward, Interstitial cells of cajal as pacemakers in the gastrointestinal tract. Annu. Rev. Physiol. 68, 307–343 (2006).

14. C. G. Richter, M. Babo-Rebelo, D. Schwartz, C. Tallon-Baudry, Phase-amplitude coupling at the organism level: The amplitude of spontaneous alpha rhythm fluctuations varies with the phase of the infra-slow gastric basal rhythm. NeuroImage 146, 951–958 (2017).

15. I. Rebollo, A.-D. Devauchelle, B. Béranger, C. Tallon-Baudry, Stomach-brain synchrony reveals a novel, delayed-connectivity resting-state network in humans. eLife 7, e33321 (2018).

16. L. Banellis, I. Rebollo, N. Nikolova, M. Allen, Stomach–brain coupling indexes a dimensional signature of mental health. Nat. Ment. Health, 1–10 (2025).

17. G. Stacher, B. Presslich, H. Stärker, Gastric acid secretion and sleep stages during natural night sleep. Gastroenterology 68, 1449–1455 (1975).

18. W. C. Orr, R. Fass, S. S. Sundaram, A. O. Scheimann, The effect of sleep on gastrointestinal functioning in common digestive diseases. Lancet Gastroenterol. Hepatol. 5, 616–624 (2020).

19. I. N. Pigarev, V. A. Bagaev, E. V. Levichkina, G. O. Fedorov, I. I. Busigina, Cortical visual areas process intestinal information during slow-wave sleep. Neurogastroenterol. Motil. 25, 268–e169 (2013).

20. I. N. Pigarev, M. L. Pigareva, The state of sleep and the current brain paradigm. Front. Syst. Neurosci. 9 (2015).

21. D. Azzalini, I. Rebollo, C. Tallon-Baudry, Visceral signals shape brain dynamics and cognition. Trends Cogn. Sci. 23, 488–509 (2019).

22. E. A. Mayer, Gut feelings: the emerging biology of gut–brain communication. Nat. Rev. Neurosci. 12, 453– 466 (2011).

23. T. Andrillon, Y. Nir, R. J. Staba, F. Ferrarelli, C. Cirelli, G. Tononi, I. Fried, Sleep spindles in humans: insights from intracranial EEG and unit recordings. J. Neurosci. 31, 17821–17834 (2011).

24. A. B. L. Tort, R. Komorowski, H. Eichenbaum, N. Kopell, Measuring phase-amplitude coupling between neuronal oscillations of different frequencies. J. Neurophysiol. 104, 1195–1210 (2010).

25. L. M. J. Fernandez, A. Lüthi, Sleep spindles: mechanisms and functions. Physiol. Rev. 100, 805–868 (2020).

26. B. P. Staresina, J. Niediek, V. Borger, R. Surges, F. Mormann, How coupled slow oscillations, spindles and ripples coordinate neuronal processing and communication during human sleep. Nat. Neurosci. 26, 1429– 1437 (2023).

27. M. Steriade, A. Nuñez, F. Amzica, A novel slow (< 1 Hz) oscillation of neocortical neurons in vivo: depolarizing and hyperpolarizing components. J. Neurosci. Off. J. Soc. Neurosci. 13, 3252–3265 (1993).

28. A. R. Adamantidis, C. Gutierrez Herrera, T. C. Gent, Oscillating circuitries in the sleeping brain. Nat. Rev. Neurosci. 20, 746–762 (2019).

29. Y. Wei, G. P. Krishnan, M. Komarov, M. Bazhenov, Differential roles of sleep spindles and sleep slow oscillations in memory consolidation. PLoS Comput. Biol. 14, e1006322 (2018).

30. T. Schreiner, M. Petzka, T. Staudigl, B. P. Staresina, Respiration modulates sleep oscillations and memory reactivation in humans. Nat. Commun. 14, 8351 (2023).

31. V. Ghibaudo, M. Juventin, N. Buonviso, L. Peter-Derex, The timing of sleep spindles is modulated by the respiratory cycle in humans. Clin. Neurophysiol. Off. J. Int. Fed. Clin. Neurophysiol. 166, 252–261 (2024).

32. J. Lechinger, D. P. J. Heib, W. Gruber, M. Schabus, W. Klimesch, Heartbeat-related EEG amplitude and phase modulations from wakefulness to deep sleep: Interactions with sleep spindles and slow oscillations. Psychophysiology 52, 1441–1450 (2015).

33. C. Mikutta, M. Wenke, K. Spiegelhalder, E. Hertenstein, J. G. Maier, C. L. Schneider, K. Fehér, J. Koenig, A. Altorfer, D. Riemann, C. Nissen, B. Feige, Co-ordination of brain and heart oscillations during non-rapid eye movement sleep. J. Sleep Res. 31, e13466 (2022).

34. Z. I. Lázár, D.-J. Dijk, A. S. Lázár, Infraslow oscillations in human sleep spindle activity. J. Neurosci. Methods 316, 22–34 (2019).

35. S. F. Schoch, M. J. Cordi, M. Schredl, B. Rasch, The effect of dream report collection and dream incorporation on memory consolidation during sleep. J. Sleep Res. 28, e12754 (2019).

36. C. da Estrela, J. McGrath, L. Booij, J.-P. Gouin, Heart rate variability, sleep quality, and depression in the context of chronic stress. Ann. Behav. Med. Publ. Soc. Behav. Med. 55, 155–164 (2020).

37. M. Naji, G. P. Krishnan, E. A. McDevitt, M. Bazhenov, S. C. Mednick, Coupling of autonomic and central events during sleep benefits declarative memory consolidation. Neurobiol. Learn. Mem. 157, 139–150 (2019).

38. N. Maingret, G. Girardeau, R. Todorova, M. Goutierre, M. Zugaro, Hippocampo-cortical coupling mediates memory consolidation during sleep. Nat. Neurosci. 19, 959–964 (2016).

39. R. Vallat, V. D. Shah, M. P. Walker, Coordinated human sleeping brainwaves map peripheral body glucose homeostasis. Cell Rep. Med. 4, 101100 (2023).

40. K. N. Browning, R. A. Travagli, Central nervous system control of gastrointestinal motility and secretion and modulation of gastrointestinal functions. Compr. Physiol. 4, 1339–1368 (2014).

41. M. Elam, P. Thorén, T. H. Svensson, Locus coeruleus neurons and sympathetic nerves: Activation by visceral afferents. Brain Res. 375, 117–125 (1986).

42. T. Bolt, S. Wang, J. S. Nomi, R. Setton, B. P. Gold, B. deB. Frederick, B. T. T. Yeo, J. J. Chen, D. Picchioni, J. H. Duyn, R. N. Spreng, S. D. Keilholz, L. Q. Uddin, C. Chang, Autonomic physiological coupling of the global fMRI signal. Nat. Neurosci. 28, 1327–1335 (2025).

43. N. E. Fultz, G. Bonmassar, K. Setsompop, R. A. Stickgold, B. R. Rosen, J. R. Polimeni, L. D. Lewis, Coupled electrophysiological, hemodynamic, and cerebrospinal fluid oscillations in human sleep. Science 366, 628– 631 (2019).

44. A. K. Seth, A. B. Barrett, L. Barnett, Granger causality analysis in neuroscience and neuroimaging. J. Neurosci. 35, 3293–3297 (2015).

45. C. J. Quinn, N. Kiyavash, T. P. Coleman, Directed information graphs. IEEE Trans. Inf. Theory 61, 6887– 6909 (2015).

46. B. Bonaz, V. Sinniger, S. Pellissier, Vagus nerve stimulation at the interface of brain–gut interactions. Cold Spring Harb. Perspect. Med. 9, a034199 (2019).

47. D. J. Levinthal, P. L. Strick, Multiple areas of the cerebral cortex influence the stomach. Proc. Natl. Acad. Sci. 117, 13078–13083 (2020).

48. A. A. Gharibans, S. Kim, D. C. Kunkel, T. P. Coleman, High-resolution electrogastrogram: a novel, noninvasive method for determining gastric slow-wave direction and speed. IEEE Trans. Biomed. Eng. 64, 807–815 (2017).

49. A. B. Allegra, A. A. Gharibans, G. E. Schamberg, D. C. Kunkel, T. P. Coleman, Bayesian inverse methods for spatiotemporal characterization of gastric electrical activity from cutaneous multi-electrode recordings. PLOS ONE 14, e0220315 (2019).

50. G. O’Grady, T. R. Angeli, P. Du, C. Lahr, W. J. E. P. Lammers, J. A. Windsor, T. L. Abell, G. Farrugia, A. J. Pullan, L. K. Cheng, Abnormal initiation and conduction of slow-wave activity in gastroparesis, defined by high-resolution electrical mapping. Gastroenterology 143, 589–598.e3 (2012).

51. S. F. Schoch, L. Salvesen, B. Rasch, M. S. Navid, S. Ataei, J. Windt, M. Schredl, M. Dresler, G. Bernardi, The relationship of memory consolidation with task incorporations into dreams – A registered report. TWCF Consciousness Studies [Preprint] (2023). 10.17605/OSF.IO/7DWJZ.

52. A. Rao, M. Dresler, I. Rebollo, J. Zeitzer, S. F. Schoch, T. Coleman, Materials associated with Rao A.A., et al. (2026), entitled “Dynamic stomach-brain electrical coupling during human sleep.”. doi: 10.25740/hy063cd6153 (2026).

53. A. T. Beck, C. H. Ward, M. Mendelson, J. Mock, J. Erbaugh, An inventory for measuring depression. Arch. Gen. Psychiatry 4, 561–571 (1961).

54. A. T. Beck, N. Epstein, G. Brown, R. A. Steer, An inventory for measuring clinical anxiety: psychometric properties. J. Consult. Clin. Psychol. 56, 893–897 (1988).

55. R. Görtelmeyer, SF-A/R Und SF-B/R: Schlaffragebogen A Und B (Hogrefe, Göttingen).

56. G. H. Klem, H. O. Lüders, H. H. Jasper, C. Elger, The ten-twenty electrode system of the International Federation. The International Federation of Clinical Neurophysiology. Electroencephalogr. Clin. Neurophysiol. Suppl. 52, 3–6 (1999).

57. N. Bigdely-Shamlo, T. Mullen, C. Kothe, K.-M. Su, K. A. Robbins, The PREP pipeline: standardized preprocessing for large-scale EEG analysis. *Front*. Neuroinformatics 9, 16 (2015).

58. M. Perslev, S. Darkner, L. Kempfner, M. Nikolic, P. J. Jennum, C. Igel, U-Sleep: resilient high-frequency sleep staging. Npj Digit. Med. 4, 72 (2021).

59. S. Leach, G. Sousouri, R. Huber, ‘High-Density-SleepCleaner’: An open-source, semi-automatic artifact removal routine tailored to high-density sleep EEG. J. Neurosci. Methods 391, 109849 (2023).

60. A. A. Gharibans, B. L. Smarr, D. C. Kunkel, L. J. Kriegsfeld, H. M. Mousa, T. P. Coleman, Artifact rejection methodology enables continuous, noninvasive measurement of gastric myoelectric activity in ambulatory subjects. Sci. Rep. 8, 5019 (2018).

61. M. J. Prerau, R. E. Brown, M. T. Bianchi, J. M. Ellenbogen, P. L. Purdon, Sleep neurophysiological dynamics through the lens of multitaper spectral analysis. Physiology 32, 60–92 (2017).

62. R. Vallat, M. P. Walker, An open-source, high-performance tool for automated sleep staging. eLife 10, e70092 (2021).

63. E. Maris, R. Oostenveld, Nonparametric statistical testing of EEG-and MEG-data. J. Neurosci. Methods 164, 177–190 (2007).

64. M. Vinck, M. van Wingerden, T. Womelsdorf, P. Fries, C. M. A. Pennartz, The pairwise phase consistency: A bias-free measure of rhythmic neuronal synchronization. NeuroImage 51, 112–122 (2010).

65. L. Brehm, P. M. Alday, Contrast coding choices in a decade of mixed models. J. Mem. Lang. 125, 104334 (2022).

