## Supplemental Material for "Dynamic stomach-brain electrical coupling in human sleep"

**The PDF file includes:**

Materials and Methods
Supplementary Text
Figs. S1 to S11
Tables S1 to S23
References (53-65)

**Other Supplementary Materials for this manuscript include the following:**

NA

### Materials and Methods

Subject enrollment and analysis subsets: The parent study enrolled 105 healthy participants, of whom 60 (44 female; mean age 23.1 ± 2.8 years; mean BMI 22.5 ± 3.9 kg/m$^2$) contributed EGG for at least one night as part of a protocol on sleep, dreaming, and memory consolidation. All participants provided written informed consent under a protocol approved by CMO Regio Arnhem-Nijmegen. Psychiatric and sleep-health screening followed the registered report (*51*) and included the Beck Depression Inventory (*53*) and Beck Anxiety Inventory (*54*); individuals with psychiatric illness, sleep disorders, medication affecting sleep-wake regulation, or heavy caffeine use were excluded.

Analyses were conducted at the level of participant-session nights with artifact-clean EEG–EGG recordings following preprocessing and EGG signal-quality control. Of the 120 planned nights, 108 were available for preprocessing (59 control, 49 experimental), as some participants discontinued after the control night. Three control nights were excluded due to insufficient EEG quality, yielding a core physiology dataset of 105 nights (56 control, 49 experimental).

Sleep-quality analyses were restricted to the control cohort to avoid confounds from experimentally induced awakenings. Within this cohort, N3 sleep-quality regressions included 54 nights for coupled SO-spindle events (due to insufficient events in two nights) and all 56 nights for uncoupled SO events and EGG/EEG ISO analyses.

In the experimental cohort, memory analyses were performed on the subset with complete behavioral data. Residualized morning recall was available for 47 nights, as two lacked paired evening-morning memory scores.

Stage- and event-based analyses (including EGG phase concentration, event-triggered EGG amplitude, and sigma-ISO phase-binning analyses) were conducted on the maximal eligible subset for each analysis. Sample sizes varied across analyses when nights lacked sufficient clean or bouts N2/N3 sleep, adequate numbers of detected events, or stage-specific data for summary measures. Exact sample sizes are reported in the corresponding result tables.

Experimental overview, sleep-quality ratings, and memory task: Participants completed a control night and an experimental night as described in the registered report (*51*). During the control night, participants slept for approximately an 8-hour sleep opportunity beginning around 23:00 h. The morning after sleep, participants completed the Schlaffragebogen-A (SF-AR) questionnaire (Table 10) (*55*). The 22 SF-AR questionnaire items were used to compute a composite subjective sleep quality score by reverse-scoring negatively valenced items, such that higher values indicated better perceived sleep quality.

During experimental nights, participants completed a word-picture association task adapted from prior work (*35*). Learning comprised three study blocks and two immediate recall blocks separated by a short break. On the following morning, participants completed task recall. One rater independently scored cued recall responses as correct when the described image matched the associated target. Memory retention was quantified as the change between evening learning performance and morning delayed recall, and residualized morning recall scores from the cleaned feature set were used for the physiology-behavior analyses.

Electrophysiological recordings: Whole-night polysomnography was recorded using a BrainVision acquisition system with a 64-channel actiCAP slim EEG cap arranged according to the international 10-20 system (*56*). Additional channels recorded electrooculography (2 channels), submental electromyography (3 channels), electrocardiography (2 channels), and electrogastrography (4 channels). All signals were sampled at 500 Hz. The EEG signal was referenced to the vertex during acquisition.

EGG was recorded from four abdominal electrodes arranged as a bipolar montage following prior electrogastrography work (Fig. 1B) (*12*). Electrodes were placed in three rows over the abdomen, with the negative derivation placed 4 cm to the left of the positive electrode. The midpoint between the xiphoid process and the umbilicus was identified, and the first electrode pair was set 2 cm below this area, with the negative derivation (1-) set at a point below the rib cage closest to the left mid-clavicular line. The second electrode pair (2+, 2-) was set 2 cm above the umbilicus and aligned with the first electrode pair. The positive derivation of the third pair (3+) was set in the center of the square formed by electrode pairs 1 and 2. The positive derivation of the fourth electrode pair (4+) was centered on the line traversing the xiphoid process and umbilicus at the same level as the third electrode. The ground electrode was placed above the iliac crest. The montage was used to derive the gastric rhythm in the canonical normogastric band centered near 0.05 Hz.

Sleep scoring and preprocessing: EEG recordings were first inspected in MATLAB to verify behavioral triggers and recording boundaries. Preprocessing was then performed in Python using MNE-Python. Continuous data were segmented from lights-off to lights-on, band-pass filtered, and trimmed to remove periods without valid recordings. Sleep staging relied on frontal, central, and occipital EEG derivations together with EOG channels, with noisy channels replaced with neighboring derivations when necessary after automated quality screening with PyPREP (*57*). Cleaned data were exported to EDF and scored with the U-Sleep algorithm using six EEG channels (two each from frontal, central and occipital regions) (*58*).

Automatic staging was validated against human scoring on a subset of nights. If epoch-wise agreement exceeded 70%, the remaining datasets were accepted with targeted manual review limited to low-confidence periods (certainty score $< 4$ for $\geq 3$ consecutive minutes). Final hypnograms consisted of 30-second epochs labeled Wake, N1, N2, N3, or REM.

EEG and EGG data then underwent semi-automatic artifact rejection using the High-Density SleepCleaner toolbox (*59*). Artifacts were marked in 10-s segments by trained raters and stored as binary masks. Noisy EEG segments were excluded or interpolated across neighboring sensors when possible, and long contaminated stretches were discarded. Clean EEG used for oscillatory analyses was band-pass filtered between 0.5 and 35 Hz.

For EGG, segments containing prolonged artifact ($\geq$ 3 minutes) were excluded. Remaining traces were notch filtered at 50 Hz and denoised with the Wiener plus adaptive-thresholding procedure described by Gharibans et al. (*60*). For each night, the EGG channel with the clearest gastric-band peak was selected for downstream analyses.

EGG signal characterization and quality control (Figs. 1; Fig. S1): To verify that the retained abdominal signal was gastric rather than residual cardiac or motion contamination, we quantified several night-level signal-quality metrics before the main physiological analyses. First, we computed each night's dominant EGG peak frequency from the abdominal power spectral density and summarized its distribution across nights. Second, we quantified the fraction of total signal power contained in the gastric band (0.03-0.07 Hz) and compared this with the fraction in the cardiac band (0.8-1.5 Hz). Third, we regressed out residual cardiac activity and measured the percent change in gastric-band power before and after this regression, together with the correlation between pre- and post-regression gastric power estimates. Finally, we computed gastric-cardiac phase-locking values (PLVs) and converted them to z-scores relative to within-night null distributions obtained by time shifting, to confirm that persistent cardiac locking did not explain the retained gastric signal.

These signal-quality analyses were used descriptively to document that the selected abdominal channel exhibited a stable gastric rhythm centered near 0.05 Hz, contained substantially more power in the gastric than the cardiac band, and was minimally altered by cardiac regression.

Gastric power across sleep cycles and sleep stages (Fig. 2): To quantify slow-wave gastric activity across the night, the selected EGG trace was downsampled as needed and analyzed with sliding-window spectral methods. To extract gastric-band power multitaper spectrograms (*61*) within the gastric frequency range, 0.03 to 0.07 Hz (*12*), filters were applied to the EGG signal using 400-s windows, 1-s step size, five tapers, and a time-bandwidth product of 2. Gastric band power was obtained by summing spectral power across the gastric frequency range at each time point. Whole-night EGG spectrograms were used to visualize stage-dependent gastric activity, and gastric power was then summarized per sleep cycle and per sleep stage.

Cycle- and stage-level gastric power effects were tested with linear mixed-effects models (LMMs) implemented in the statsmodels Python package. Models included fixed effects for sleep cycle or sleep stage, experimental condition, sex, and time since last meal, with random intercepts for participant. These analyses generated the main Fig. 2 cycle trajectory as well as the stage contrasts and their supplementary breakdowns.

Extraction of EGG infraslow oscillations (ISOs) (Fig.2; Fig. S2): To isolate slower modulation of gastric power itself, we computed a multitaper spectrogram (*61*) of the gastric-band power time series using 300-s windows, 1-s step size, five tapers, and time-bandwidth product of 3, focusing on 0.005-0.1 Hz. This yielded an EGG power ISO spectrum for each night and sleep stage. Stage-dependent EGG ISO power was then quantified in the 0.006-0.010 Hz range and compared across NREM, REM, and wake using LMMs with the same covariate structure described above.

Detection of cortical slow oscillations and spindles: Slow oscillations (SOs) and spindles were detected from clean EEG using YASA (*62*). Spindles were detected in all clean EEG channels between 10 to 16 Hz (*23*) and were selected if: 1) their duration lasted between 0.5 and 2 seconds, 2) the minimum distance between two spindles was 500 milliseconds, and 3) fit the default relative power, moving correlation, and root mean square detection thresholds (*62*). SOs were detected in all clean EEG channels between 0.5 and 4 Hz (*6*) and were selected if: 1) the minimum and maximum duration of the negative deflection fell between 0.3 and 1.5 seconds, 2) the duration of the positive deflection fell between 0.1 and 1 second, 3) amplitude of the absolute negative trough fell between 40 and 200$\mu$V, 4) amplitude of the absolute positive peak fell between 10 and 150$\mu$V, and 5) amplitude of the peak to peak fell between 75 and 350$\mu$V (*62*). Event detection was restricted to N2 and N3 sleep.

An SO was classified as spindle-coupled when a spindle peak occurred within 1.5 seconds after the SO negative peak, consistent with prior work on SO-spindle grouping during the up-state rebound (*26*). Events without such a spindle were treated as uncoupled. Similarly, a spindle was classified as coupled if its peak occurred within 1.5 seconds after the SO negative peak (i.e., during the SO up-state).

<u>EEG amplitude modulation by gastric phase during NREM events (Fig. 3B; Fig. S4):</u> To quantify gastric phase-amplitude coupling (PAC) between the stomach rhythm and cortical activity, the EGG signal was band-pass filtered in the gastric band (0.03-0.07 Hz) and its instantaneous phase was extracted with the Hilbert transform. For each EEG channel, narrow-band amplitude envelopes were computed from 0.5 and 25.5 Hz in 1-Hz steps. Modulation index (MI) (*24*) was then calculated from 20 phase bins spanning 0 to $2\pi$ during artifact-free N2 and N3 data segments.

At each frequency, phases were binned into 20 equal intervals between 0 and 2π, and the mean amplitude within each bin was normalized to form a phase-amplitude distribution. MI was computed as the Kullback-Leibler divergence from a uniform distribution (*24*). Within-night surrogate distributions were generated by circularly time-shifting one signal relative to the other (60 seconds, corresponding to ~ 3 EGG cycles for gastric phase) and recomputing MI across 400 shuffles. For each channel and frequency, a signed z-score was computed as $Z = \frac{MI_{emp} - \mu_{sur}}{\sigma_{sur}}$ where $\mu_{sur}$ and $\sigma_{sur}$ were the surrogate mean and standard deviation, respectively.

For the scalp maps shown in the main figure, MI values were averaged within the delta (0.5-4 Hz (*6*)) and sigma (10-16 Hz (*23*)) ranges and tested against zero with one-sample cluster based permutation tests across channels (10,000 permutations; cluster-forming threshold set at the analytical t-critical value at $\alpha$ = 0.05). Channels with clusters $p$ < 0.05 were considered significant (FWER controlled) and highlighted on the topomap (*63*). The supplementary heatmaps were produced from the full frequency-by-channel MI matrices for N2 and N3 (10,000 permutations; cluster-forming threshold set at the analytical t-critical value at $\alpha$ = 0.05) (*63*).

<u>Gastric phase concentration during SOs and SO-spindle complexes (Fig. 3D-E; Fig. S5):</u> Instantaneous EGG phase was sampled at the SO trough and, for coupled events, at the spindle peak occurring after the trough. For each participant, stage, coupling condition, and region of interest, the gastric phase distribution was modeled as a von Mises distribution and estimated with maximum likelihood using the scipy.stats.vonmises Python package. We summarized these distributions with the von Mises concentration parameter $\kappa$, which reflects the sharpness of the peak of the probability density (*64*). Fits required at least five events, and uncoupled events were count-matched to the number of coupled events with repeated resampling when necessary. These $\kappa$ values were analyzed with mixed-effects models that tested coupling effects, stage effects, and ROI effects, together with cycle-level changes across the night in the supplementary analyses. In

the main figure, paired coupled-versus-uncoupled comparisons were emphasized for N2 and N3, with ROI-specific contrasts reported in the supplements.

Event-triggered gastric amplitude modulation around SO troughs (Fig. 3F, G; Fig. S6): To test whether coupled cortical events were accompanied by transient changes in gastric amplitude, EGG was filtered in the gastric band and converted to analytic amplitude with the Hilbert transform. For each retained SO event, we extracted ±40 s around the SO negative peak and z-scored the amplitude relative to the pre-event baseline (-40 to -10 s). Within each night, coupled and uncoupled traces were averaged separately.

For scalar summaries, we compared the average amplitude in pre-event (−20 to 0 s) and post-event (0 to 20 s) windows and computed the post-minus-pre change for coupled and uncoupled events. These values, reflecting baseline-normalized EGG amplitude change, are reported in the main figure (Fig. 3F, G). Coupled and uncoupled curves were compared both as time series and as summary change scores, with the main figure reporting the NREM average trace and the stage-specific distributions (Fig. S6A).

To assess whether these changes exceeded chance expectations, we further normalized post–pre differences relative to within-night surrogate distributions generated by randomly jittering SO event timing within ±20 s while requiring the full analysis window to remain within the same sleep stage and preserving event counts. This yielded z-scored values of (observed − $\text{mean}_{\text{surrogate}}$) / $(\text{standard deviation})_{\text{surrogate}}$, where positive scores indicate greater modulation than expected by chance. These null-normalized values are reported in supplementary analyses (Fig. S6B).

Amplitude changes were analyzed with mixed-effects models that tested coupling effects, stage effects, and ROI effects in the supplementary analyses. In the main figure, paired coupled-versus uncoupled comparisons were emphasized for N2 and N3, with ROI-specific contrasts reported in the supplements (Fig. S6C, D).

Extraction of EEG sigma infraslow oscillations (Fig. 4; Fig. S7): To characterize infraslow oscillations (ISOs) in the EEG sigma power band, short-time multitaper spectrograms (*61*) were computed for each EEG channel in the sigma band (10-16 Hz) using 4-s windows, 1-s step size, five tapers, and time-bandwidth product of 3. Sigma-band power was summed across the sigma range to yield a sigma-power time series sampled at 10 Hz. This time series was used both for stage-level sigma power analyses and for slower infraslow modulation analyses.

A second spectral decomposition (*61*) was then applied to the sigma-power time series to extract EEG sigma-power ISOs. Sigma ISO power was summarized by sleep stage, and the prominence of the canonical ~0.02 Hz sigma ISO peak was quantified in NREM sleep. These analyses correspond to the sigma-stage and sigma-ISO panels in the supplementary figure.

Spatiotemporal coupling between EEG sigma ISOs and EGG power ISOs (Fig. 4B, C; Fig. S8): To align EGG and EEG sigma ISOs, the EEG sigma-power time series and gastric power time series were interpolated to the same sigma-derived time base and low-pass filtered with a 4th-order Butterworth filter (0.2 Hz cutoff). Because gastric power was estimated from much longer windows (300 s) than sigma power, the gastric series was shifted to align the centers of the two analysis windows.

Both signals were segmented into non-overlapping 5-minute bouts within each sleep stage. Within each bout, the signals were z-scored and cross-correlated across symmetric time lags. Bout-level cross-correlograms were averaged within channel and then across participants to obtain stage-level spatiotemporal maps. Statistical significance of stage-specific cross-correlation clusters was assessed with non-parametric cluster-based permutation testing in MNE, using adjacency across both sensors and lag samples. Clusters were formed using a cluster-forming t-threshold corresponding to $p = 0.05$ (two-tailed), and cluster-level significance was evaluated with 10,000 permutations at $\alpha = 0.01$ (two-tailed, cluster-level corrected).

<u>EGG ISO phase modulation of EEG sigma ISO amplitude (Fig. 4D; Fig. S9):</u> We next quantified PAC between the phase of the EGG power ISO and the amplitude of EEG sigma-power ISO. The EGG ISO phase was derived from the 0.005-0.009 Hz range, while EEG sigma ISO amplitude was evaluated across 0.01-0.10 Hz frequencies. Analytic signals were obtained with the Hilbert transform and MI was calculated as above from continuous N2 and N3 bouts of at least 5 minutes. Night-level MI values were converted to z-scores relative to circular-shift (180 seconds, which is ~1.5 EGG ISO cycles, for the EGG ISO phase) surrogate distributions as described above (*24*). To determine stage-dependent variations in EGG and EEG sigma ISO, we computed the difference in z-scored MI values between N2 and N3 within EEG sigma ISO frequencies (0.02 to 0.1 Hz) per channel. We ran a cluster-based permutation test (10,000 permutations; cluster-forming threshold set by the analytical t-critical value at $\alpha = 0.05$, two-sided). Channels belonging to clusters with $p < 0.05$ (FWER controlled) were highlighted on a topomap in the main figure (*63*). The supplementary heatmaps were produced from the full frequency-by-channel MI matrices for N2 and N3 (10,000 permutations; cluster-forming threshold set at the analytical t-critical value at $\alpha = 0.05$) (*63*).

<u>Phase-binned sigma-state, spindle-probability, and gastric-amplitude analyses (Fig. 4E; Fig. S10):</u> To test whether cortical and gastric event dynamics followed the phase of the sigma ISO, valid sigma-ISO cycles were defined from consecutive zero crossings in the low-pass-filtered sigma power signal, requiring cycle durations of 25 to 100 s. Each cycle was divided into eight ordered phase bins (0 to $2\pi$) spanning the descending phase, trough, ascending phase, and peak. For each subject, values were averaged within phase bins across cycles to obtain subject-level estimates of sigma power (z-scored), spindle probability, and EGG power. Samples not assigned to a valid sigma ISO cycle formed the "No ISOs" control, excluding samples outside the bounds of accepted cycles within each stage segment. Phase modulation was assessed using mixed-effects models with sinusoidal ($\sin(\phi)$, $\cos(\phi)$) terms and permutation-based tests (1,000 permutations). Planned paired contrasts compared trough (bins 1-4), peak (bins 5-8), and No-ISO periods, with Holm correction applied across contrasts.

<u>Memory recall association analyses (Fig. 5A, Fig. S11A-B; Table S20):</u> Behavioral relevance for memory was tested in experimental-session nights with complete physiology and behavioral data ($n = 47$ nights). An initial exploratory screen evaluated the candidate physiology features listed in Fig. S11A and Table 11 using one-at-a-time partial correlations with Benjamini–Hochberg false

discovery rate correction applied within the screen. Based on prior hypotheses regarding stage-specific stomach–brain coordination during NREM sleep, confirmatory analyses focused on two sigma-band coupling metrics shown in Fig. 5A: (i) EEG sigma-EGG PAC during N2 sleep (N2 coupling) and (ii) the stage-dependent EEG sigma-EGG PAC difference between N3 and N2 sleep (N3–N2 coupling). Memory and predictor values were residualized on sex, time since last meal, N2 and N3 SO-spindle coupling percentage, and N2 and N3 spindle density using Huber robust regression, and associations were summarized using partial Spearman correlations.

Subjective sleep quality and intrinsic physiology analyses (Fig. 5B, C; Table 13; Fig. S11C, D): Sleep-quality analyses were restricted to control nights (n = 56 nights). An initial exploratory screen evaluated candidate physiology features against objective and subjective sleep quality measures (Fig. S11C) using one-at-a-time partial correlations with Benjamini–Hochberg false discovery rate correction within the screen. Confirmatory analyses focused on gastric phase concentration during N3 sleep and intrinsic infraslow variability measures based on their physiological relevance to NREM arousability and gastric dynamics. For Fig. 5B, gastric phase concentration values were related to subjective sleep quality after residualizing for canonical sleep architecture (percentage of N1, N2, and N3 sleep; sleep-to-wake transitions per hour; and N3-to-N1/N2 transitions per hour), together with sex and time since last meal. Huber robust regression was used, and partial Spearman correlations yielded the regression lines shown in Fig. 5B. For Fig. 5C, subjective sleep quality was similarly residualized on canonical sleep architecture and related to intrinsic EGG ISO variability and EEG sigma ISO variability. The same adjusted framework was used for the comparison between EGG ISO variability and HRV RMSSD, allowing gastric and cardiac predictors to be evaluated within the same residualized space (Fig. S11D).

Statistical framework: The paper combines several complementary statistical approaches matched to the structure of each analysis. Cycle- and stage-level gastric analyses and several event-level physiology summaries were tested with linear mixed-effects models in the statsmodels Python package. These models included the analysis-relevant fixed effects of sleep stage or sleep cycle, experimental condition where applicable, sex, and time since last meal, together with random intercepts for repeated observations within participant or night as appropriate to the dataset being analyzed. Continuous predictors were mean-centered and categorical predictors were contrast-coded (*64*) where noted.

All mixed-effects models were fit using restricted maximum likelihood (REML) estimation with the L-BFGS optimizer, and significance was assessed using two-sided Wald tests on the fixed effect coefficients. Standardized effect sizes were reported as model-based standardized coefficients or model-estimated mean differences with 95% confidence intervals. Multiple comparisons were controlled within analysis families using the Benjamini-Hochberg false-discovery rate (FDR; $q < 0.05$), except for the planned trough/peak/No-ISO paired contrasts in Fig. 4E, which used Holm correction.

Spatial EEG analyses and spatiotemporal cross-correlograms were evaluated with cluster-based permutation tests in MNE-Python. Behavioral regression figures were evaluated using Huber robust regression and partial Spearman correlations. The same covariate-adjustment framework was maintained across analyses to ensure comparability across predictors.

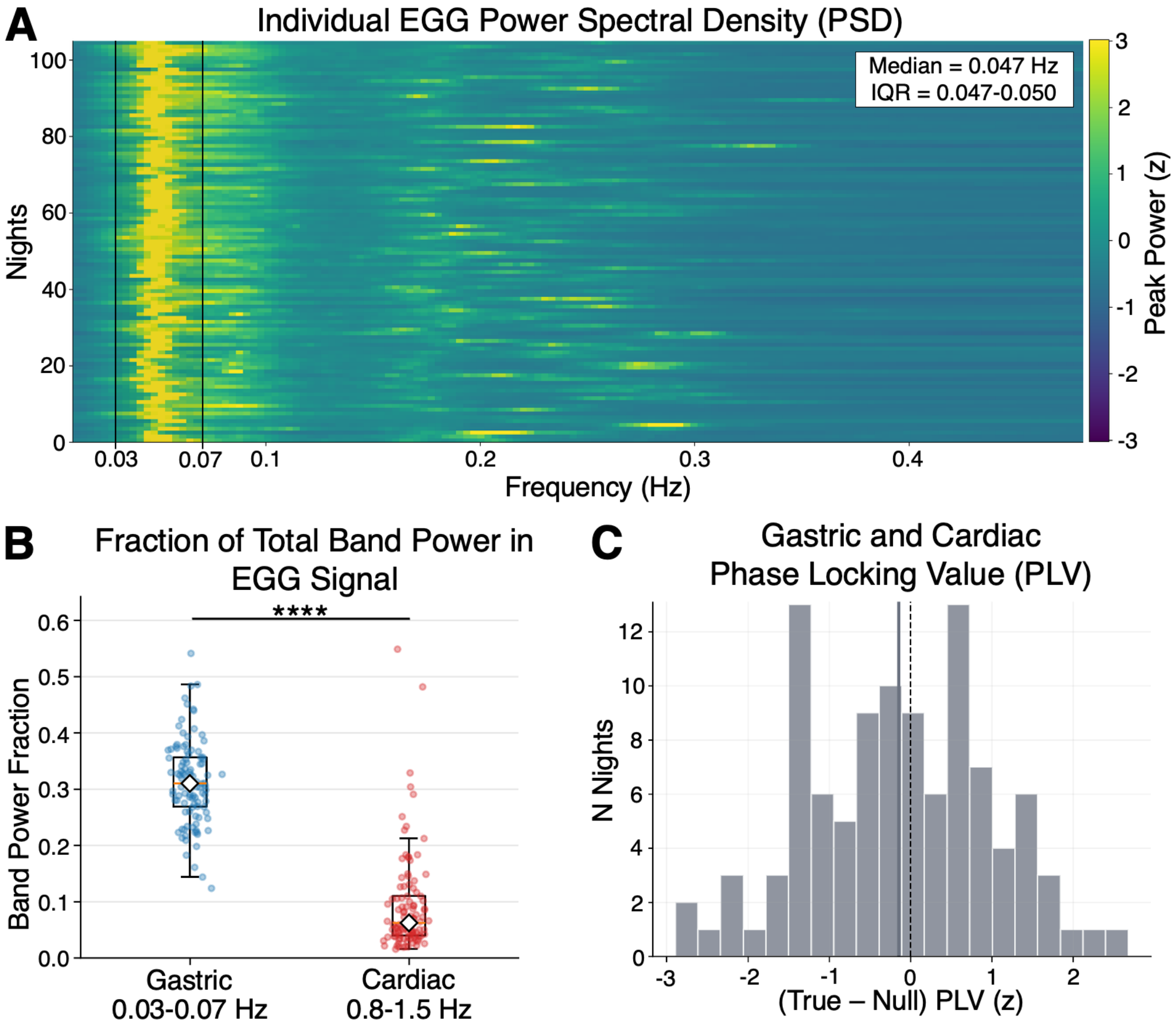


**Fig. S1**. **Gastric rhythms dominate EGG spectral power and are independent of cardiac activity. (A)** Individual power spectral densities (PSDs) of the EGG signal (0.02-0.5 Hz) across all nights show a prominent peak within the normogastric frequency band (0.03-0.07 Hz; black box). **(B)** The gastric frequency band constitutes a significantly larger fraction of total power (0.01-1.5 Hz) compared to the cardiac frequency band (0.8-1.5 Hz). **(C)** Phase-locking between gastric and ECG-derived cardiac signals does not exceed the null distribution, indicating no significant coupling (n = 104 nights).

ns (not significant) $p \geq 0.05$, * $p < 0.05$, ** $p < 0.01$, *** $p < 0.001$, **** $p < 0.0001$. All values corrected for multiple comparisons when necessary. See Tables S1-S2 for statistical details.

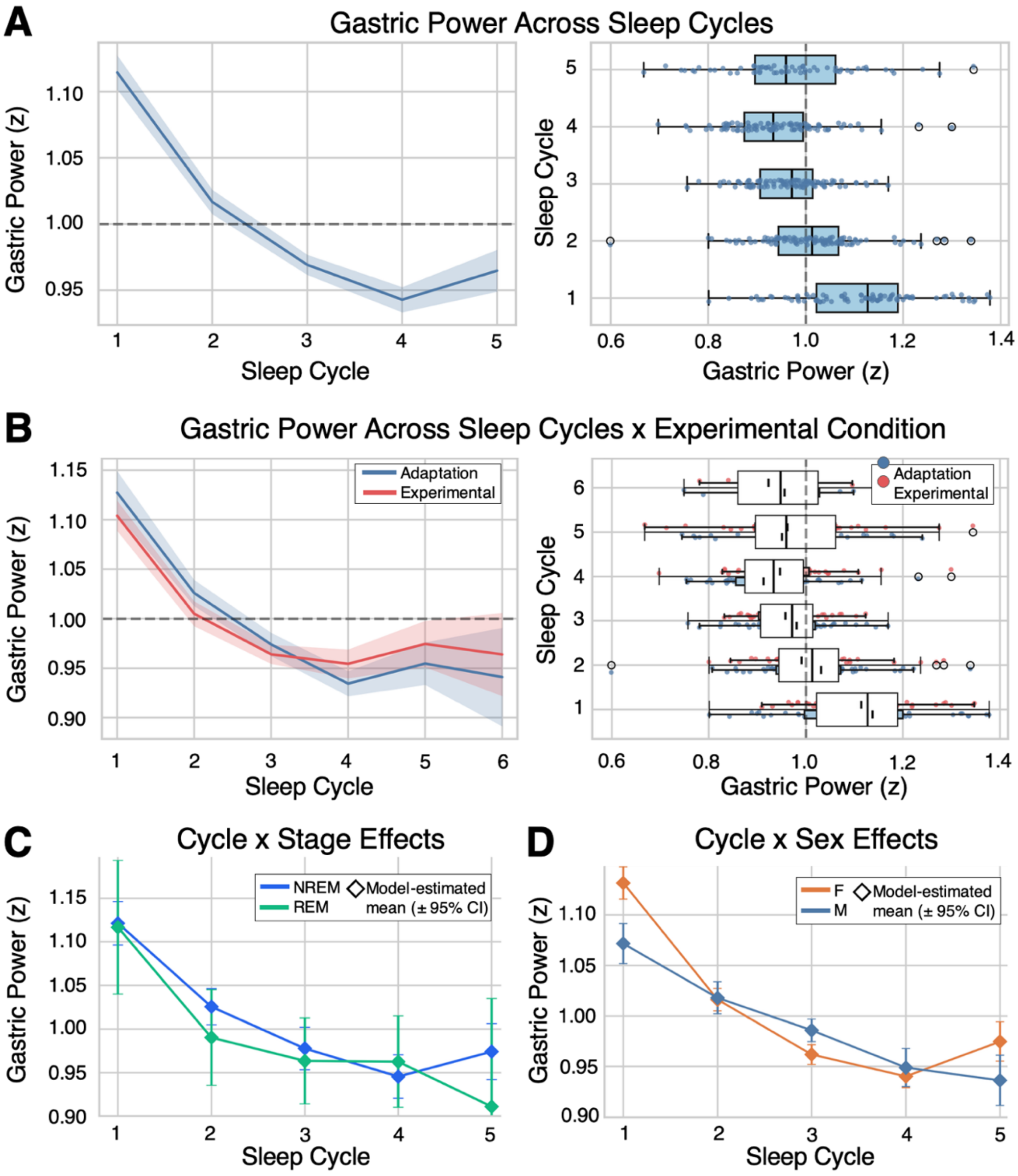


**Fig. S2**. **Gastric power declines across sleep cycles with stage- and sex-dependent modulation but no effect of condition. (A)** Gastric power decreases across successive sleep cycles. **(B)** This decline is consistent across control and experimental nights, with no significant main effect of condition. **(C)** Gastric power shows a comparable stage-by-cycle interaction with similar declining trajectories across NREM and REM sleep. **(D)** Gastric power did not exhibit a significant sex-by-cycle interaction. (n = 105 nights). See Tables 1, S3, and S4 for statistical details and sample sizes.

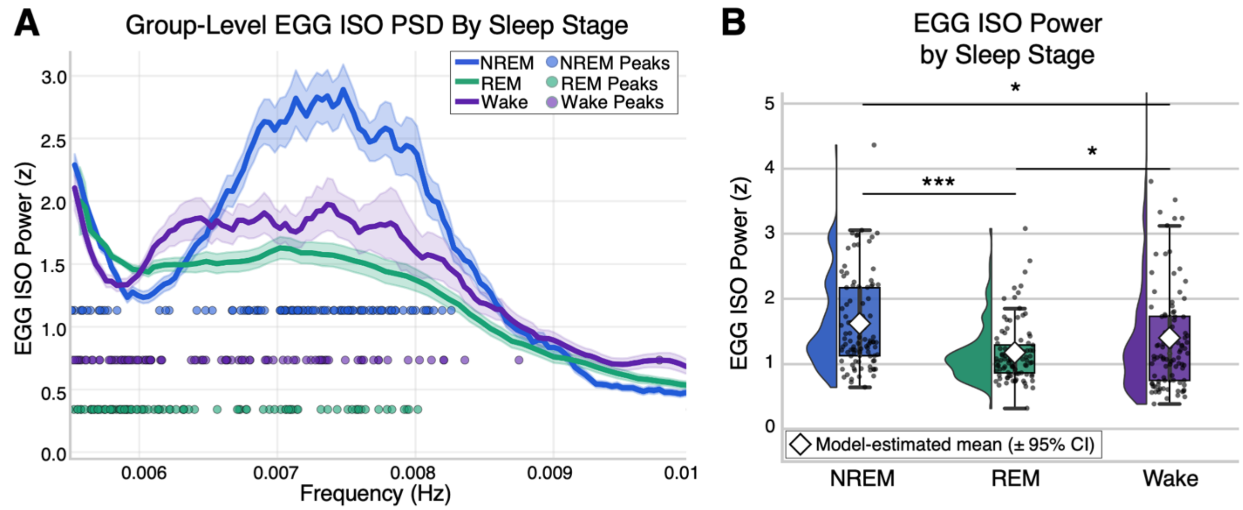


**Fig. S3**. **Gastric infraslow oscillations are strongest during NREM sleep relative to REM and wake. (A)** Group-level EGG ISO power spectral density (PSD) across sleep stages (NREM, REM, wake) shows a clear infraslow peak ( 0.007 Hz), with greatest power during NREM sleep. Shaded regions denote ± SEM, and dots indicate individual peak frequencies. **(B)** EGG ISO power is significantly higher in NREM compared to REM and wake, with no differences between REM and wake. Adjusted means represent estimated marginal means (± 95% CI) from linear mixed-effects models (n = 99 nights).

ns (not significant) $p \geq 0.05$, * $p < 0.05$, ** $p < 0.01$, *** $p < 0.001$, **** $p < 0.0001$. All values corrected for multiple comparisons when necessary. See Table S5 for statistical details and sample sizes.

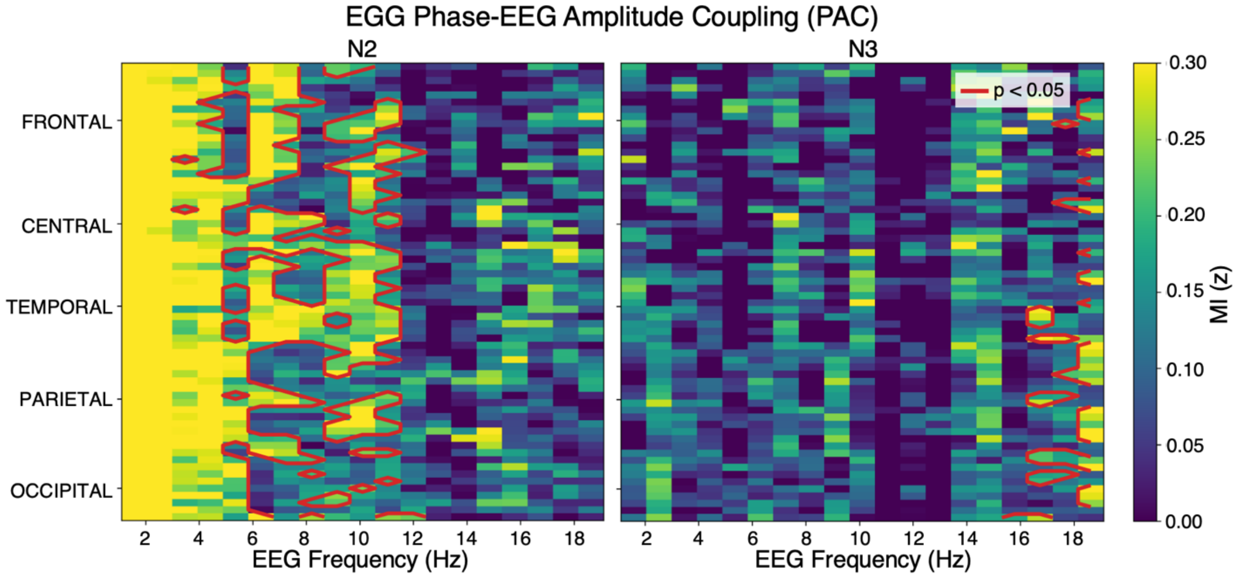


**Fig. S4**. **EGG-EEG phase-amplitude coupling is prominent in N2 but absent in N3 sleep.** Channel-by-frequency EEG-EGG phase-amplitude coupling (PAC) heatmaps across EEG regions in N2 (left) and N3 (right) sleep. Warmer colors indicate stronger modulation index (modulation index (MI), z-scored). In N2, significant PAC clusters are observed across low-frequency bands and widespread cortical regions (outlined in red; cluster-corrected $p < 0.05$, n = 105 nights). In contrast, no significant PAC clusters are detected in N3 (n = 105 nights).

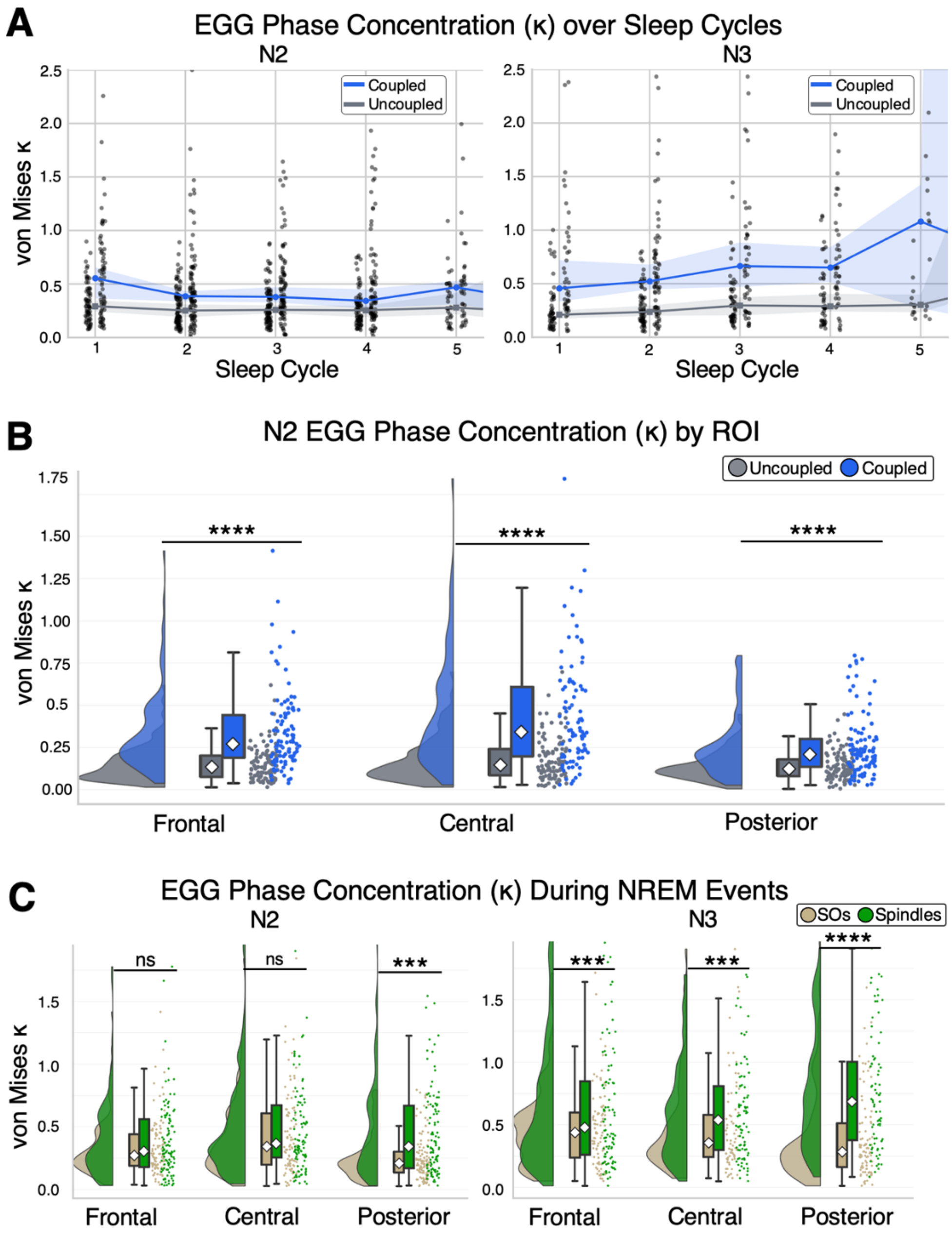


**Fig. S5**. **Gastric phase concentration increases with SO-spindle coupling and shows stage-, cycle-, and region-specific effects. (A)** EGG phase concentration ($\kappa$) across sleep cycles in N2 (left) and N3 (right), shown separately for coupled and uncoupled events. No significant cycle effects are observed in N2, whereas N3 shows a significant increase in $\kappa$ across cycles, independent of coupling. **(B)** In N2, $\kappa$ is significantly higher during coupled compared to uncoupled SO events across all regions of interest (frontal, central, posterior). **(C)** When SO-spindle coupling occurred, EGG $\kappa$ increases occurred during spindle peaks rather than SO troughs. This effect was present in posterior regions during N2 (left) and all regions during N3 (right).

ns (not significant) $p \geq 0.05$, * $p < 0.05$, ** $p < 0.01$, *** $p < 0.001$, **** $p < 0.0001$. All values corrected for multiple comparisons when necessary. See Table S6-9 for statistical details and sample sizes.

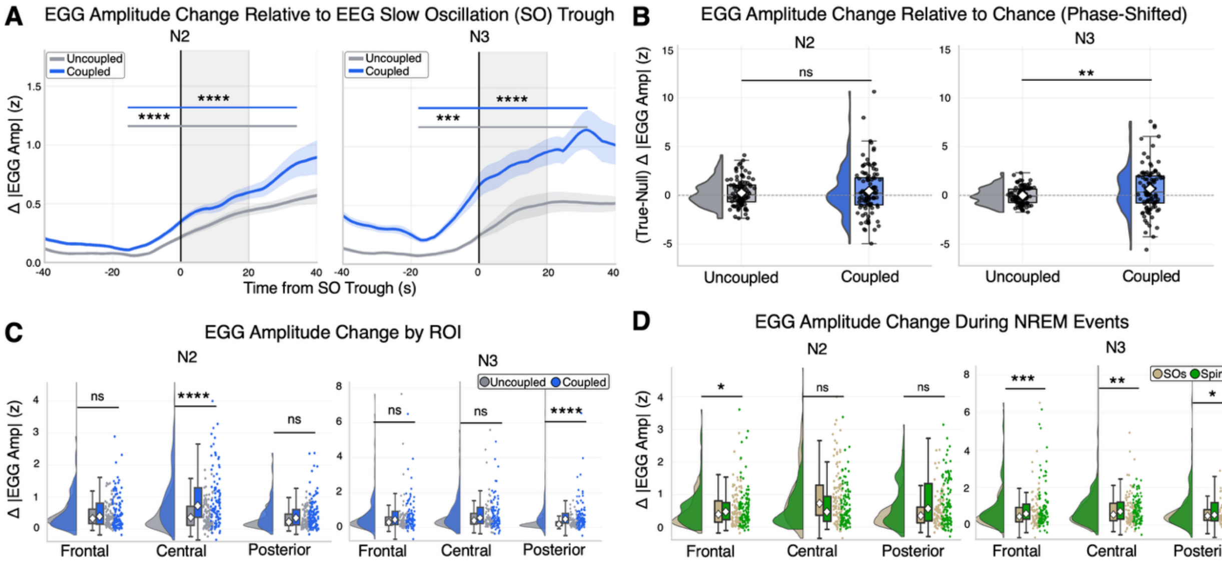


**Fig. S6**. **SO-locked gastric amplitude increases after the trough and is enhanced by SO-spindle coupling. (A)** Time course of baseline-referenced absolute EGG amplitude aligned to EEG slow oscillation (SO) troughs in N2 (left) and N3 (right), shown separately for coupled and uncoupled events. Both conditions exhibit significant post-trough increases in EGG amplitude, with larger effects during coupled events. **(B)** Observed EGG amplitude changes were normalized to within-night surrogate distributions generated by shifting SO event timing, yielding values where positive scores indicate greater modulation than expected by changes. Plots demonstrate that EGG amplitude changes during coupled SO-spindle events are greater than chance-level during N3 (right) than N2 (left). **(C)** In N2, EGG amplitude is significantly higher during coupled SO events than uncoupled events across central regions. Whereas during N3, EGG amplitude is significantly higher during coupled SO events across posterior regions. **(D)** When SO-spindle coupling occurred, EGG amplitude increases occurred during spindle peaks rather than SO troughs. This effect was present in frontal regions during N2 (left) and frontal and central regions during N3 (right).

ns (not significant) $p \geq 0.05$, * $p < 0.05$, ** $p < 0.01$, *** $p < 0.001$, **** $p < 0.0001$. All values corrected for multiple comparisons when necessary. See Tables S10- S15 for statistical details and sample sizes.

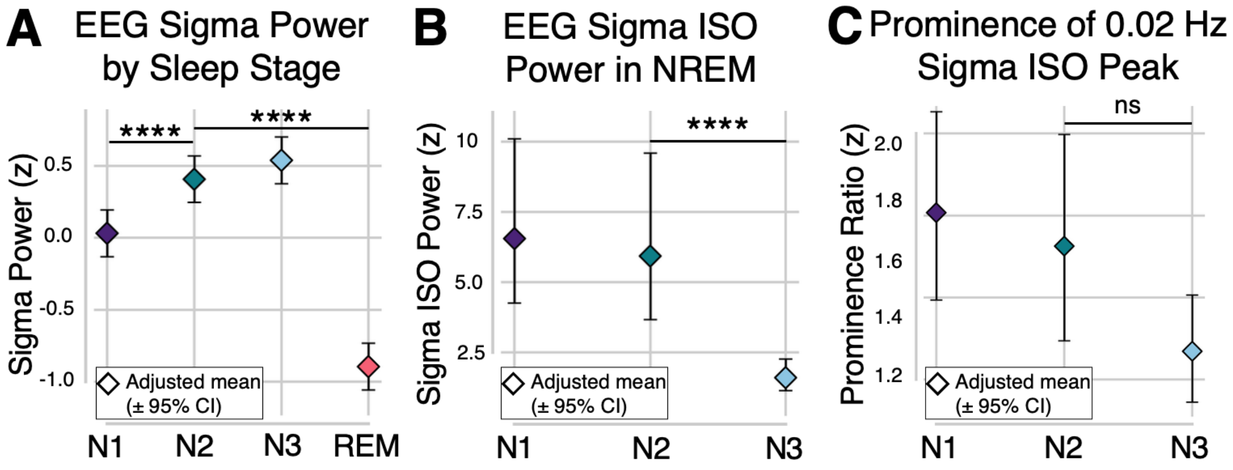


**Fig. S7**. **EEG sigma infraslow dynamics persist across NREM sleep despite reduced magnitude in N3. (A)** EEG sigma-band power varies across sleep stages, with highest levels in N2 and N3 and markedly reduced power in REM sleep. **(B)** Narrowband sigma infraslow oscillation (ISO) power (0.02 Hz) is significantly reduced in N3 compared to N2, while remaining present across NREM sleep. **(C)** The relative prominence of the 0.02 Hz sigma ISO peak is comparable in all sub-stages of NREM sleep, indicating preservation of infraslow structure. Adjusted means represent estimated marginal means ± 95% CI from covariate-adjusted linear mixed-effects models (n = 105 nights).

ns (not significant) $p \geq 0.05$, * $p < 0.05$, ** $p < 0.01$, *** $p < 0.001$, **** $p < 0.0001$. All values corrected for multiple comparisons when necessary. See Tables S16-S18 for statistical details and sample sizes.

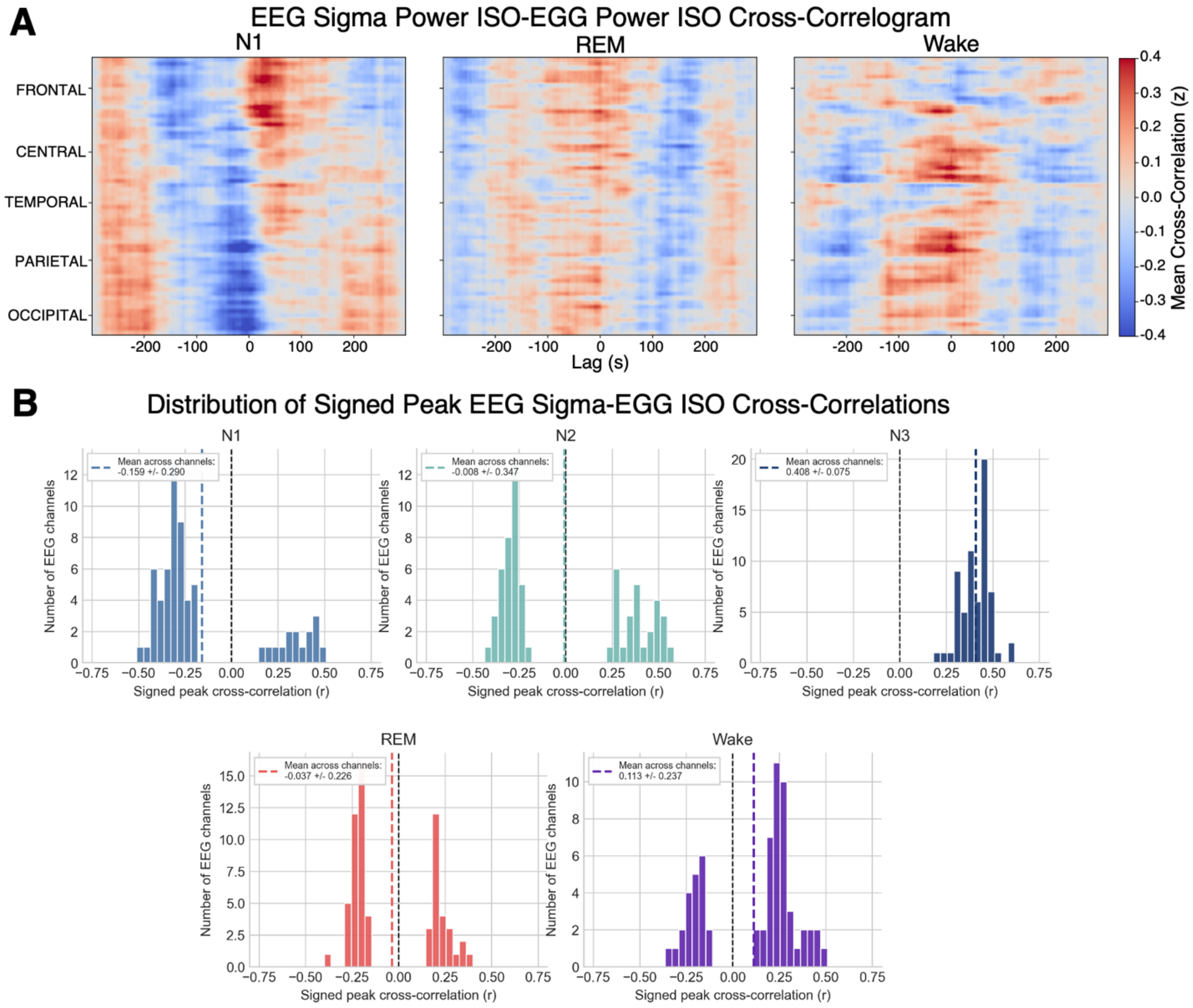


**Fig. S8**. **EEG sigma-gastric infraslow coupling reverses sign across sleep stages and is maximally positive in N3. (A)** Channel-by-lag cross-correlograms between EEG sigma power ISOs and EGG power ISOs across sleep stages (N1, REM, wake), showing heterogeneous and state-dependent coupling patterns across cortical regions (n = 105 nights). Spatiotemporal permutation cluster tests revealed no significant coupling between the two signals. **(B)** Distribution of signed peak cross-correlations across EEG channels for each sleep stage. Peak correlations were defined as the maximum absolute cross-correlation across lags, retaining sign. N1 and REM show predominantly negative correlations, N2 shows a mixed distribution, and N3 shows uniformly positive correlations across all channels, indicating a strong and consistent coupling direction. Reported values are mean ± s.e.m. across EEG channels (n = 105 nights).

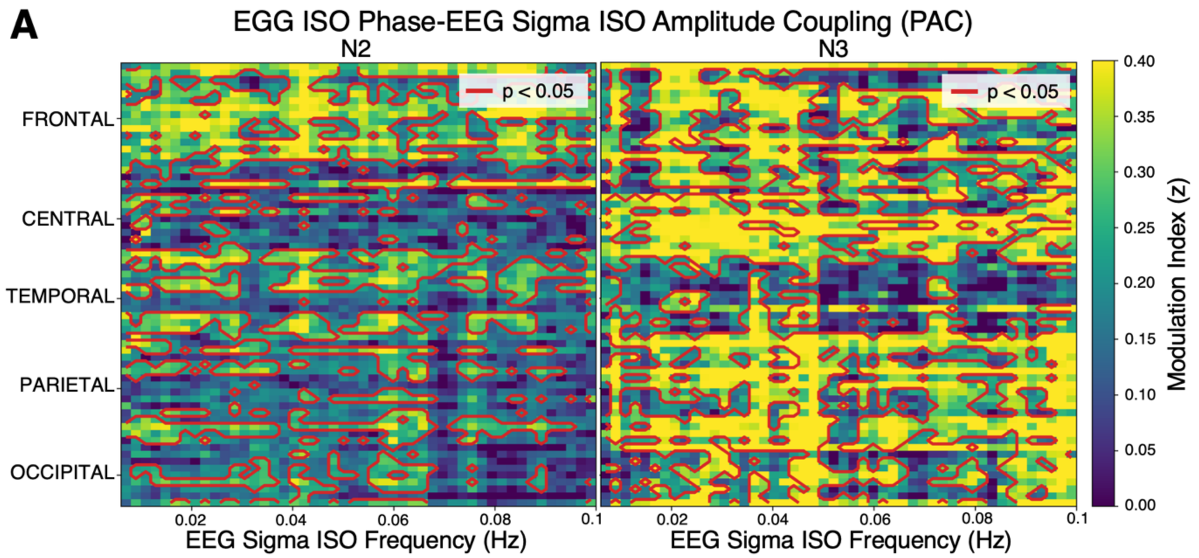


**Fig. S9**. **Cross-frequency infraslow coupling between gastric and cortical sigma dynamics is enhanced in N3 sleep. (A)** Channel-by-frequency phase-amplitude coupling (PAC) between EGG ISO phase and EEG sigma ISO amplitude in N2 (left) and N3 (right). Warmer colors indicate stronger modulation index (MI, z-scored). Significant positive clusters are observed in both stages (red contours; cluster-corrected $p < 0.05$), with more widespread and stronger coupling in N3 ($n = 105$ nights).

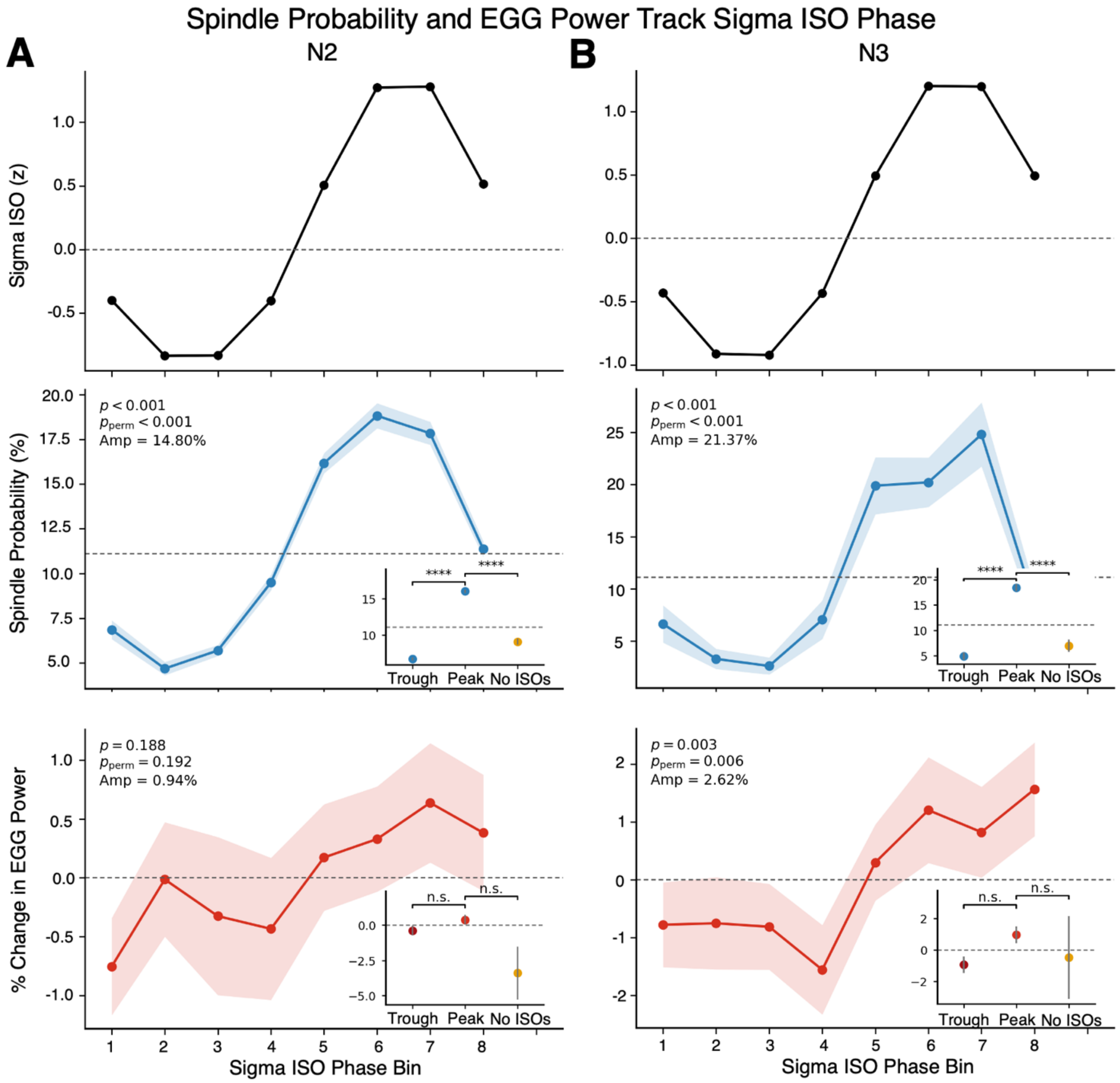


**Fig. S10. Spindle probability and gastric activity are modulated by sigma infraslow phase across NREM sleep. (A)** In N2 sleep, spindle probability (middle) varies systematically across the phase of the sigma infraslow oscillation (top), with maximal values near the peak phase, while EGG power change (bottom) does not show significant phase modulation. **(B)** In N3 sleep, spindle probability exhibits strong phase modulation across the sigma infraslow cycle, while EGG power shows a smaller but statistically significant modulation pattern. Trough values summarize lower-phase bins (1 - 4), peak values summarize higher-phase bins (5 - 8), and no-ISO values summarize samples outside detected sigma ISOs. Omnibus p-values are from mixed-effects models with sin(ϕ) and cos(ϕ) terms fit across bins, and permutation p -values are based on 1000 within-unit phase shuffles (n = 103 nights).

ns (not significant) $p \geq 0.05$, * $p < 0.05$, ** $p < 0.01$, *** $p < 0.001$, **** $p < 0.0001$. All values corrected for multiple comparisons when necessary. See Table S19 for statistical details and sample sizes.

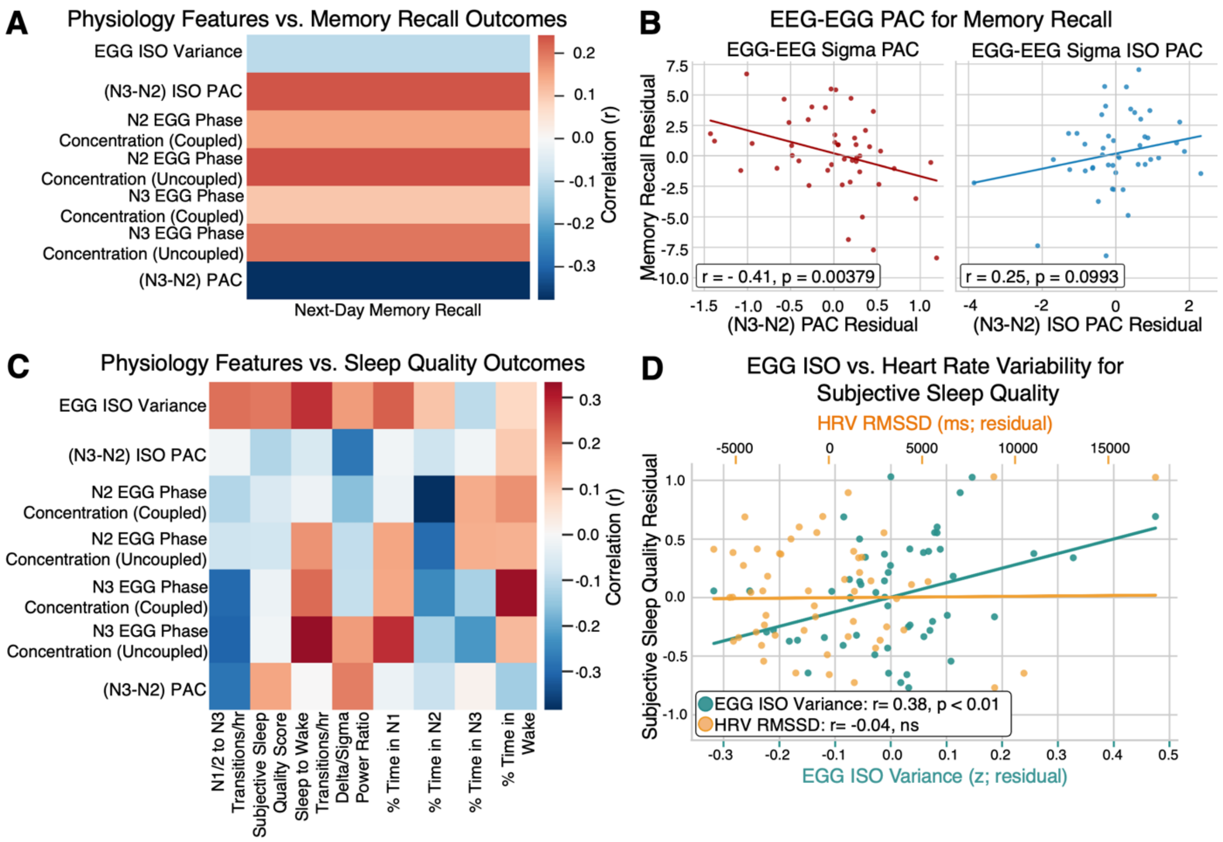


**Fig. S11**. **Gastric infraslow dynamics and uncoupled N3 phase organization relate to subjective sleep quality. (A)** Exploratory associations between physiological features and memory recall outcomes during experimental nights (n = 47 nights). Each feature was tested independently against next-day memory recall after residualizing sex and time since last meal. **(B)** Increased N3-N2 EEG sigma-EGG PAC (red; r = -0.41, p < 0.01) was significantly associated with worse next day memory recall (left), whereas increased N3-N2 EEG sigma-EGG ISO PAC had a positive trend with improved memory recall (blue; r = 0.25, p = ns; n = 47 nights). Memory recall scores were residualized on EEG-derived measures supportive of memory consolidation such as SO-spindle coupling strength and spindle density. **(C)** Exploratory associations between physiological features and sleep outcomes during control nights (n = 56 nights). Each feature was tested independently against subjective sleep quality after residualizing sex and time since last meal. **(D)** Comparison of gastric and cardiac predictors of subjective sleep quality (n = 56 nights). Residualized subjective sleep quality is positively associated with EGG ISO variability (teal; r = 0.38, p < 0.01), whereas no significant relationship is observed with heart rate variability (HRV RMSSD; yellow; r = -0.04, p = ns).

All residualizations were performed using Huber robust regression, and partial Spearman correlations were used for visualization and summary statistics. Reported values are partial correlations. See Tables S20-23 for statistical details and sample sizes.

**Table S1**. **Whole-night gastric-band and cardiac-band power fractions.** Values are mean ± s.e.m. across nights presented in Fig. S1B. The statistical test is a paired t-test comparing gastric-band and cardiac-band fractions of total power across nights.

| Band | # nights | Mean ± s.e.m. | Test statistic | p-value |
|---|---|---|---|---|
| **Gastric (0.03–0.07 Hz)** | 105 | 0.312 ± 0.007 | $t(104) = 16.220$ | $3.03 \times 10^{-30}$ |
| **Cardiac (0.8–1.5 Hz)** | 105 | 0.092 ± 0.009 | | |

**Table S2. Gastric-cardiac phase-locking relative to the null distribution.** The first panel summarizes the observed PLV relative to the null 95th percentile from Fig. S1C. The second panel reports the null-standardized PLV z-score summary from the same nights.

| Comparison | # nights | Observed PLV mean ± s.e.m. | Null95 mean ± s.e.m. | Summary |
|---|---|---|---|---|
| **Gastric ↔ Cardiac** | 104 | 0.000098 ± 0.000024 | 0.000126 ± 0.000026 | 5/104 exceed null 95th (p= 0.5991) |

| Comparison | # nights | Mean z-score | Median z-score | Statistical test |
|---|---|---|---|---|
| **Gastric ↔ Cardiac** | 104 | -0.138 | -0.142 | One-sample t-test vs 0 (t = -1.242, p = 0.8914) |

**Table S3**. **Stage-by-cycle interaction in hourly gastric power.** Mixed-effects model is shown for the plotted cycles from Fig. S2C. The omnibus interaction was tested by likelihood-ratio comparison between mixed-effects models with and without the interaction term.

| Term | Coef. | s.e. | z | p-value | 95% CI |
|---|---|---|---|---|---|
| **Intercept** | 1.1251 | 0.0141 | 79.774 | < 0.0001 | [1.0975, 1.1528] |
| **REM sleep cycle** | -0.0085 | 0.0405 | -0.210 | 0.8339 | [-0.0880, 0.0710] |
| **Sleep cycle 2** | -0.1004 | 0.0148 | -6.785 | < 0.0001 | [-0.1293, -0.0714] |
| **Sleep cycle 3** | -0.1487 | 0.0160 | -9.267 | < 0.0001 | [-0.1802, -0.1173] |
| **Sleep cycle 4** | -0.1799 | 0.0163 | -11.027 | < 0.0001 | [-0.2118, -0.1479] |
| **Sleep cycle 5** | -0.1516 | 0.0191 | -7.936 | < 0.0001 | [-0.1891, -0.1142] |
| **Condition (Control vs Exp)** | -0.0078 | 0.0102 | -0.765 | 0.4442 | [-0.0278, 0.0122] |
| **Sex (M vs F)** | -0.0079 | 0.0109 | -0.720 | 0.4715 | [-0.0293, 0.0135] |
| **REM × Cycle 2** | -0.0262 | 0.0502 | -0.522 | 0.6019 | [-0.1246, 0.0722] |
| **REM × Cycle 3** | -0.0102 | 0.0488 | -0.209 | 0.8348 | [-0.1058, 0.0855] |
| **REM × Cycle 4** | 0.0256 | 0.0500 | 0.511 | 0.6096 | [-0.0725, 0.1236] |
| **REM × Cycle 5** | -0.0544 | 0.0780 | -0.697 | 0.4857 | [-0.2072, 0.0985] |
| **Time since meal (h)** | 0.0000 | 0.0015 | 0.031 | 0.9756 | [-0.0028, 0.0029] |

| Test | $\chi^2$ | df | p-value |
|---|---|---|---|
| **Sleep cycle × Sleep Stage (NREM vs REM)** | 2.24 | 4 | 0.6920 |

**Table S4**. **Sex-by-cycle interaction in hourly gastric power.** Mixed-effects model is shown for the plotted cycles and sexes from Fig. S2D. The omnibus interaction was tested by likelihood-ratio comparison between mixed-effects models with and without the interaction term.

| Term | Coef. | s.e. | z | p-value | 95% CI |
|---|---|---|---|---|---|
| **Intercept** | 1.1372 | 0.0152 | 74.969 | < 0.0001 | [1.1074, 1.1669] |
| **Sleep cycle 2** | -0.1208 | 0.0165 | -7.300 | < 0.0001 | [-0.1533, -0.0884] |
| **Sleep cycle 3** | -0.1751 | 0.0186 | -9.426 | < 0.0001 | [-0.2115, -0.1387] |
| **Sleep cycle 4** | -0.1939 | 0.0192 | -10.093 | < 0.0001 | [-0.2316, -0.1562] |
| **Sleep cycle 5** | -0.1587 | 0.0230 | -6.905 | < 0.0001 | [-0.2038, -0.1137] |
| **Sex (M vs F)** | -0.0641 | 0.0243 | -2.640 | 0.0083 | [-0.1116, -0.0165] |
| **Condition (Control vs Exp)** | -0.0049 | 0.0102 | -0.486 | 0.6272 | [-0.0248, 0.0150] |
| **Cycle 2 × Sex** | 0.0674 | 0.0318 | 2.119 | 0.0341 | [0.0050, 0.1297] |
| **Cycle 3 × Sex** | 0.0913 | 0.0328 | 2.785 | 0.0054 | [0.0270, 0.1556] |
| **Cycle 4 × Sex** | 0.0741 | 0.0338 | 2.189 | 0.0286 | [0.0077, 0.1404] |
| **Cycle 5 × Sex** | 0.0272 | 0.0418 | 0.652 | 0.5147 | [-0.0547, 0.1092] |
| **Time since meal (centered)** | -0.0007 | 0.0014 | -0.521 | 0.6024 | [-0.0034, 0.0020] |

| Test | $\chi^2$ | df | p-value |
|---|---|---|---|
| **Sex × sleep cycle interaction** | 9.58 | 4 | 0.0482 |

**Table S5**. **NREM, REM, and Wake EGG infraslow oscillation (ISO) power.** Estimates are from the linear mixed model output presented in Fig. S3B. Estimated marginal means are reported for each stage, alongside pairwise LMM contrasts with FDR-corrected p-values.

| Stage | # nights | Estimated marginal mean | Statistical test |
|---|---|---|---|
| **NREM** | 99 | 1.5940 [1.5862, 1.6020] | Linear mixed model |
| **REM** | 99 | 1.3479 [1.3401, 1.3559] | Linear mixed model |
| **Wake** | 99 | 1.3052 [1.2974, 1.3132] | Linear mixed model |

| Comparison | Effect size (β) | Test stat. (z) | Corrected p-value | $d_{LMM}$ |
|---|---|---|---|---|
| **NREM vs REM** | 0.4455 | 4.657 | 0.0000 | 0.683 |
| **NREM vs Wake** | 0.2192 | 2.267 | 0.0116 | 0.336 |
| **REM vs Wake** | 0.2263 | 2.198 | 0.0279 | 0.347 |

**Table S6**. **Sleep-cycle counts for EGG phase concentration analyses.** Per-cycle night counts contributing to the sleep-cycle panel from Fig. S5A.

| Stage | Sleep cycle | # nights uncoupled | # nights coupled |
|---|---|---|---|
| **N2** | 1 | 61 | 60 |
| **N2** | 2 | 90 | 90 |
| **N2** | 3 | 85 | 85 |
| **N2** | 4 | 66 | 66 |
| **N2** | 5 | 25 | 25 |
| **N2** | 6 | 3 | 3 |
| **N3** | 1 | 53 | 53 |
| **N3** | 2 | 59 | 59 |
| **N3** | 3 | 36 | 35 |
| **N3** | 4 | 22 | 21 |
| **N3** | 5 | 9 | 7 |
| **N3** | 6 | 2 | 2 |

**Table S7**. **Sleep-cycle terms from the stage-wise EGG phase concentration LMMs.** Results from fitted stage-wise covariate-adjusted linear mixed-effects models on log($\kappa$) with coupling, sleep cycle, coupling × sleep cycle, condition, time since meal, and sex as predictors presented in Fig. S5A. Holm p-values correct across N2 and N3 within each reported model term.

| **Stage** | **Model term** | **# nights** | **β ± s.e.** | **Test statistic** | **Raw p** | **Holm p** |
|---|---|---|---|---|---|---|
| **N2** | Sleep cycle | 104 | 0.002 ± 0.024 | z = 0.066 | 0.9478 | 0.9478 |
| **N2** | Coupling × cycle | 104 | -0.023 ± 0.022 | z = -1.035 | 0.3005 | 0.3005 |
| **N3** | Sleep cycle | 92 | 0.091 ± 0.040 | z = 2.272 | 0.0231 | 0.0462 |
| **N3** | Coupling × cycle | 92 | -0.089 ± 0.035 | z = -2.528 | 0.0115 | 0.0229 |

**Table S8**. **ROI comparison of coupled versus uncoupled SO events in N2.** EGG phase concentration ($\kappa$) by ROI during SO events in N2, comparing coupled and uncoupled events. Displayed values in Fig. S5B are per-night median [IQR] of raw $\kappa$. Statistical tests are simple effects from a covariate-adjusted linear mixed-effects model fit on log($\kappa$) with ROI, sleep stage, coupling, condition, sex, and time since meal as predictors.

| ROI | (# nights) Median uncoupled [95% CI] | (# nights) Median coupled [95% CI] | β ± s.e. | Test stat. | Corrected p-value |
|---|---|---|---|---|---|
| **Frontal** | (105) 0.14 [0.07, 0.20] | (99) 0.26 [0.15, 0.43] | 0.721 ± 0.108 | z = 6.643 | < 1e-10 |
| **Central** | (105) 0.14 [0.09, 0.22] | (89) 0.39 [0.20, 0.78] | 1.029 ± 0.112 | z = 9.205 | < 1e-10 |
| **Posterior** | (105) 0.12 [0.07, 0.18] | (103) 0.23 [0.16, 0.40] | 0.821 ± 0.107 | z = 7.692 | < 1e-10 |

| ROI | Fold change |
|---|---|
| **Frontal** | × 2.06 |
| **Central** | × 2.80 |
| **Posterior** | × 2.27 |

**Table S9**. **LMM effects for coupled spindle versus coupled SO events within ROI.** Estimates are simple effects for coupled spindle − coupled SO on log($\kappa$) within each ROI, fit separately for N2 and N3 and presented in Fig. S5C. Because only coupled events are analyzed here, the coupling term is constant in usable data and is dropped from the model formula. Ratios are exp($\beta$). FDR q-values correct across ROIs within stage.

| Stage | ROI | SO nights | Spindle nights | β ± s.e. | Test stat. (z) | Raw p | FDR q | Ratio |
|---|---|---|---|---|---|---|---|---|
| **N2** | Frontal | 99 | 102 | 0.294 ± 0.124 | 2.375 | 0.0176 | 0.0263 | × 1.342 |
| **N2** | Central | 89 | 94 | 0.186 ± 0.128 | 1.451 | 0.1467 | 0.1467 | × 1.205 |
| **N2** | Posterior | 103 | 102 | 0.365 ± 0.122 | 2.995 | 0.0027 | 0.0082 | × 1.441 |
| **N3** | Frontal | 93 | 77 | 0.488 ± 0.131 | 3.728 | 0.0002 | 0.0006 | × 1.630 |
| **N3** | Central | 65 | 83 | 0.146 ± 0.143 | 1.018 | 0.3085 | 0.3085 | × 1.157 |
| **N3** | Posterior | 92 | 88 | 0.266 ± 0.127 | 2.096 | 0.0361 | 0.0542 | × 1.305 |

**Table S10**. **Stage-specific within-stage pre-post comparisons of EGG amplitude.** Displayed values in Fig. S6A are raw night-level mean ± s.e.m. in the prespecified analysis windows (Pre: −40 to −20 s; Post: 0 to 20 s) computed from baseline-referenced absolute EGG amplitude. Statistical tests were within-coupling pre-post contrasts extracted from stage-specific linear mixed-effects models with coupling condition, experimental condition, sex, and time since last meal as fixed effects, and subject/night repeated measures.

| Stage | Coupling | Pre mean ± s.e.m. | Post mean ± s.e.m. | Est. diff. | Test stat. (z) | p-val | Effect size ($d_{LMM}$) |
|---|---|---|---|---|---|---|---|
| **N2** | Uncoupled | 0.051 ± 0.003 | 0.364 ± 0.033 | 0.313 | 8.265 | < 0.0001 | 1.141 |
| **N2** | Coupled | 0.095 ± 0.006 | 0.545 ± 0.047 | 0.450 | 11.885 | < 0.0001 | 1.640 |
| **N3** | Uncoupled | 0.058 ± 0.008 | 0.408 ± 0.068 | 0.350 | 3.886 | 0.0001 | 0.541 |
| **N3** | Coupled | 0.171 ± 0.016 | 0.967 ± 0.130 | 0.796 | 8.847 | < 0.0001 | 1.233 |

**Table S11**. **Stage-specific null-normalization analysis of event-triggered EGG amplitude.** Displayed values in Fig. S6B are raw night-level mean ± s.e.m. of the null-normalized post-pre change score ($z\Delta$) from the mixed-model analysis subset for coupled and uncoupled events. The primary statistical test was the stage-specific Coupled−Uncoupled contrast from a linear mixed-effects model with event count, experimental condition, sex, time since last meal, and normalized delta power as covariates, with repeated measures accounted for by subject and night random effects.

| Stage | Uncoupled mean ± s.e.m. | Coupled mean ± s.e.m. | Estimated diff. | Test stat. (z) | p-val | Effect size ($d_{LMM}$) |
|---|---|---|---|---|---|---|
| **N2** | 0.291 ± 0.132 | 0.471 ± 0.238 | 0.180 | 0.703 | 0.4818 | 0.098 |
| **N3** | -0.032 ± 0.085 | 0.656 ± 0.231 | 0.688 | 2.664 | 0.0077 | 0.373 |

**Table S12**. **ROI comparison of coupled versus uncoupled SO events in N2 for EGG amplitude change.** Estimates in Fig. S6B are simple effects from the ROI-resolved mixed model fit in the ROI amplitude section. Positive estimates indicate larger Δ|EGG amplitude| for coupled than uncoupled SO events within ROI. FDR *q*-values correct across ROIs within stage.

| ROI | Estimated diff. ± s.e. | Test stat. (z) | Raw p | FDR q | Effect size |
|---|---|---|---|---|---|
| **Frontal** | -0.009 ± 0.105 | -0.084 | 0.933338 | 0.933338 | -0.012 |
| **Central** | 0.452 ± 0.106 | 4.253 | 0.000021 | 0.000063 | 0.599 |
| **Posterior** | 0.179 ± 0.105 | 1.708 | 0.087568 | 0.131352 | 0.237 |

**Table S13. ROI comparison of coupled versus uncoupled SO events in N3 for EGG amplitude change.** Estimates from Fig. S6C are simple effects from the ROI-resolved mixed model fit in the ROI amplitude section. Positive estimates indicate larger Δ|EGG amplitude| for coupled than uncoupled SO events within ROI. FDR q-values correct across ROIs within stage.

| ROI | Estimated diff. ± s.e. | Test stat. (z) | Raw p | FDR q | Effect size $d_{LMM}$ |
|---|---|---|---|---|---|
| **Frontal** | 0.194 ± 0.107 | 1.813 | 0.069895 | 0.098103 | 0.257 |
| **Central** | 0.182 ± 0.110 | 1.654 | 0.098103 | 0.098103 | 0.241 |
| **Posterior** | 0.493 ± 0.106 | 4.641 | 0.000003 | 0.000010 | 0.653 |

**Table S14. LMM effects for coupled spindle versus coupled SO events in N2 for EGG amplitude change.** Estimates in Fig. S6D are simple effects from the ROI-resolved mixed model fit in ROI amplitude. Positive estimates indicate larger Δ|EGG amplitude| for coupled spindle than coupled SO events within ROI. FDR q-values correct across ROIs within stage.

| ROI | Estimated diff. ± s.e. | Test stat. (z) | Raw p | FDR q | Effect size |
|---|---|---|---|---|---|
| Frontal | 0.315 ± 0.118 | 2.668 | 0.007626 | 0.022878 | $d_{LMM} = 0.379$ |
| Central | 0.175 ± 0.129 | 1.363 | 0.172725 | 0.259088 | $d_{LMM} = 0.211$ |
| Posterior | −0.081 ± 0.121 | −0.673 | 0.501191 | 0.501191 | $d_{LMM} = 0.098$ |

**Table S15. LMM effects for coupled spindle versus coupled SO events in N3 for EGG amplitude change.** Estimates in Fig. S6D are simple effects from the ROI-resolved mixed model fit in ROI amplitude. Positive estimates indicate larger Δ|EGG amplitude| for coupled spindle than coupled SO events within ROI. FDR q-values correct across ROIs within stage.

| ROI | Estimated diff. ± s.e. | Test stat. (z) | Raw p | FDR q | Effect size |
|---|---|---|---|---|---|
| Frontal | 0.449 ± 0.121 | 3.718 | 0.000201 | 0.000603 | $d_{LMM} = 0.541$ |
| Central | 0.388 ± 0.129 | 3.018 | 0.002547 | 0.003821 | $d_{LMM} = 0.467$ |
| Posterior | −0.237 ± 0.124 | −1.914 | 0.055562 | 0.055562 | $d_{LMM} = -0.286$ |

**Table S16**. **Stage-dependent EEG sigma-band activity.** Sigma-band activity by sleep stage (N1, N2, N3, REM) from Fig. S7A. Estimates are from a covariate-adjusted linear mixed-effects model with fixed effects for stage, condition, sex, and centered time since meal, and a random intercept for night. Adjusted means are estimated marginal means with 95% confidence intervals.

| Stage | # nights | Adjusted mean [95% CI] | Statistical test |
|---|---|---|---|
| **N1** | 105 | 0.032 [-0.130, 0.194] | Linear mixed model |
| **N2** | 105 | 0.408 [0.245, 0.570] | |
| **N3** | 105 | 0.539 [0.377, 0.701] | |
| **REM** | 103 | -0.895 [-1.059, -0.732] | |

| Comparison | Effect β [95% CI] | Test statistic | p-value | Effect Size |
|---|---|---|---|---|
| **N1 vs. N2** | -0.272 [-0.576, -0.175] | z = -3.675 | 0.0002 | $d_{LMM}$ = -0.507 |
| **N3 vs. N2** | 0.137 [-0.069, -0.332] | z = 1.286 | 0.1985 | $d_{LMM}$ = 0.177 |
| **REM vs. N2** | -1.291 [-1.504, -1.102] | z = -12.682 | < 0.0001 | $d_{LMM}$ = 1.759 |

**Table S17. Stage-dependent narrowband EEG sigma ISO power.** Narrowband EEG sigma ISO power at 0.018-0.022 Hz by sleep stage (N1, N2, N3) from Fig. S7B. Estimates are from a covariate adjusted linear mixed-effects model with fixed effects for stage, condition, sex, centered time since meal, and centered stage duration, and a random intercept for night. Adjusted means are back transformed estimated marginal means with 95% confidence intervals.

| Stage | # nights | Adjusted mean [95% CI] | Statistical test |
|---|---|---|---|
| **N1** | 102 | 6.551 [4.251, 10.097] | Linear mixed model |
| **N2** | 105 | 5.929 [3.665, 9.592] | |
| **N3** | 105 | 1.602 [1.135, 2.260] | |

| Comparison | Effect β [95% CI] | Test statistic | p-value | Effect Size | Fold Change |
|---|---|---|---|---|---|
| **N1 vs. N2** | -0.272 [-0.576, -0.175] | z = 0.286 | 0.7745 | $d_{LMM} = 0.099$ | 1.105 x |
| **N3 vs. N2** | 0.137 [-0.069, -0.332] | z = -4.911 | < 0.0001 | $d_{LMM} = -1.303$ | 0.270 x |

**Table S18**. **Stage-dependent EEG sigma ISO peak prominence.** Relative prominence of the ~0.02 Hz sigma ISO peak, defined as mean power in 0.018-0.022 Hz divided by mean power in 0.03-0.06 Hz, by sleep stage (N1, N2, N3) presented in Fig. S7C. Estimates are from a covariate-adjusted linear mixed effects model with fixed effects for stage, condition, sex, centered time since meal, and centered stage duration, and a random intercept for night. Adjusted means are back-transformed estimated marginal means with 95% confidence intervals.

| Stage | # nights | Adjusted mean [95% CI] | Statistical test |
|---|---|---|---|
| **N1** | 102 | 1.607 [1.394, 1.853] | Linear mixed model |
| **N2** | 105 | 1.525 [1.294, 1.798] | |
| **N3** | 105 | 1.269 [1.145, 1.407] | |

| Comparison | Effect β [95% CI] | Test statistic | p-value | Effect Size | Fold Change |
|---|---|---|---|---|---|
| **N1 vs. N2** | 0.023 [-0.092, 0.137] | z = 0.388 | 0.6984 | $d_{LMM} = 0.123$ | 1.054 x |
| **N3 vs. N2** | -0.080 [-0.169,0.009] | z = -1.759 | 0.0785 | $d_{LMM} = -0.433$ | 0.832 x |

**Table S19. Stage-stratified sigma ISO phase modulation.** Stage-stratified shared sigma ISO phase modulation of spindle probability and EGG power presented in Fig. S10. Trough values summarize bins 1-4, peak values summarize bins 5-8, and no-ISO values summarize samples outside detected sigma ISOs. Omnibus phase-modulation p-values are from mixed-effects models with sin($\phi$) and cos($\phi$) terms fit across bins 1-8. Permutation p-values are based on 1000 within-unit phase shuffles.

| Stage | Outcome | # session-subject units | Trough mean ± s.e.m. | Peak mean ± s.e.m. | No ISOs mean ± s.e.m. |
|---|---|---|---|---|---|
| N2 | Spindle probability (%) | 98 | 6.68 ± 0.24 | 16.05 ± 0.23 | 9.07 ± 0.44 |
| N2 | EGG power change (%) | 102 | −0.38 ± 0.36 | 0.38 ± 0.36 | −3.40 ± 1.88 |
| N3 | Spindle probability (%) | 71 | 4.89 ± 0.69 | 18.37 ± 0.71 | 6.97 ± 1.22 |
| N3 | EGG power change (%) | 101 | −0.95 ± 0.53 | 0.95 ± 0.53 | −0.49 ± 2.63 |

| Stage | Outcome | Omnibus p-value | Permutation p-value | Peak-to-trough amp. (%) | Statistical test |
|---|---|---|---|---|---|
| N2 | Spindle probability (%) | $2.49 \times 10^{-180}$ | < 0.001 | 14.80 | Mixed model, sin($\phi$) and cos($\phi$) terms |
| N2 | EGG power change (%) | 0.188 | 0.192 | 0.94 | Mixed model, sin($\phi$) and cos($\phi$) terms |
| N3 | Spindle probability (%) | $1.53 \times 10^{-24}$ | < 0.001 | 21.37 | Mixed model, sin($\phi$) and cos($\phi$) terms |
| N3 | EGG power change (%) | 0.003 | 0.006 | 2.62 | Mixed model, sin($\phi$) and cos($\phi$) terms |

| Stage | Outcome | Comparison | Holm-corrected p-val |
|---|---|---|---|
| N2 | Spindle probability (%) | Peak vs. Trough | $6.52 \times 10^{-37}$ |
| N2 | Spindle probability (%) | Peak vs. No ISOs | $1.01 \times 10^{-22}$ |
| N2 | EGG power change (%) | Peak vs. Trough | 0.288 |
| N2 | EGG power change (%) | Peak vs. No ISOs | 0.106 |
| N3 | Spindle probability (%) | Peak vs. Trough | $1.65 \times 10^{-14}$ |
| N3 | Spindle probability (%) | Peak vs. No ISOs | $4.26 \times 10^{-10}$ |
| N3 | EGG power change (%) | Peak vs. Trough | 0.146 |
| N3 | EGG power change (%) | Peak vs. No ISOs | 0.570 |

**Table S20**. **Exploratory memory screen with stage-specific N2/N3 architecture adjustment.** Predictors in Fig. S11A were tested one at a time against morning memory recall after residualizing sex, time since meal, N2 and N3 slow-oscillation-spindle coupling, and N2 and N3 spindle density. Reported corrected q-values are Benjamini-Hochberg FDR-adjusted within the stage-specific exploratory memory screen.

| Outcome | Predictor | # nights | Partial r | p-value | FDR q-value |
|---|---|---|---|---|---|
| Morning recall residual | (N3−N2) EEG sigma-EGG PAC | 47 | −0.378 | 0.0089 | 0.0620 |
| Morning recall residual | N2 uncoupled $\kappa$ | 47 | 0.243 | 0.1004 | 0.2668 |
| Morning recall residual | EEG sigma-EGG ISO PAC | 46 | 0.236 | 0.1143 | 0.2668 |
| Morning recall residual | N3 uncoupled $\kappa$ | 47 | 0.200 | 0.1779 | 0.3113 |
| Morning recall residual | N2 coupled $\kappa$ | 47 | 0.153 | 0.3057 | 0.4280 |
| Morning recall residual | N3 coupled $\kappa$ | 47 | 0.102 | 0.4940 | 0.4981 |
| Morning recall residual | EGG ISO var | 47 | −0.101 | 0.4981 | 0.4981 |

:

**Table S21. Stage-specific memory-coupling sensitivity analysis.** Memory behavioral analysis in Fig. S11B was residualized on sex, time since meal, stage-specific N2/N3 slow-oscillation-spindle coupling, and stage-specific N2/N3 spindle density. Reported correlations are partial Spearman ρ in the adjusted space and inferential p-values are from the corresponding Huber-robust regressions.

| Predictor | # nights | Partial ρ | p-value |
|---|---|---|---|
| **(N3-N2) EEG sigma-EGG PAC** | 47 | -0.414 | 0.0038 |
| **(N3-N2) EEG sigma-EGG ISO PAC** | 46 | 0.246 | 0.0993 |

**Table S22. Exploratory sleep quality screen underlying the control-night.** Visualization in Fig. S11C. Each physiology feature was tested one at a time against subjective sleep quality after residualizing sex and time since meal. Reported corrected q-values are the global Benjamini-Hochberg FDR values.

| Outcome | Predictor | # nights | Partial r | p-value | Corrected q-value |
|---|---|---|---|---|---|
| % Time in N2 | N2 coupled $\kappa$ | 56 | −0.384 | 0.0035 | 0.1970 |
| Sleep to Wake Trans/hr | N3 uncoupled $\kappa$ | 56 | 0.334 | 0.0119 | 0.2378 |
| % Time in Wake | N3 coupled $\kappa$ | 54 | 0.328 | 0.0154 | 0.2378 |
| N1/2 to N3 Trans/hr | N3 uncoupled $\kappa$ | 56 | −0.307 | 0.0213 | 0.2378 |
| % Time in N2 | N2 uncoupled $\kappa$ | 56 | −0.295 | 0.0273 | 0.2378 |
| N1/2 to N3 Trans/hr | N3 coupled $\kappa$ | 54 | −0.297 | 0.0291 | 0.2378 |
| % Time in N1 | N3 uncoupled $\kappa$ | 56 | 0.283 | 0.0345 | 0.2378 |
| Sleep to Wake Trans/hr | EGG ISO var | 56 | 0.279 | 0.0372 | 0.2378 |
| N1/2 to N3 Trans/hr | (N3-N2) EEG-EGG PAC | 56 | −0.278 | 0.0382 | 0.2378 |
| Delta/Sigma power ratio | (N3-N2) EEG-EGG ISO PAC | 52 | −0.275 | 0.0484 | 0.2710 |
| % Time in N2 | N3 coupled $\kappa$ | 54 | −0.241 | 0.0791 | 0.4027 |
| % Time in N1 | EGG ISO var | 56 | 0.229 | 0.0900 | 0.4200 |
| % Time in N3 | N3 uncoupled $\kappa$ | 56 | −0.222 | 0.1002 | 0.4318 |
| Sleep to Wake Trans/hr | N3 coupled $\kappa$ | 54 | 0.215 | 0.1190 | 0.4634 |
| N1/2 to N3 Trans/hr | EGG ISO var | 56 | 0.208 | 0.1241 | 0.4634 |
| Subjective Sleep Quality | EGG ISO var | 56 | 0.201 | 0.1372 | 0.4800 |
| Delta/Sigma power ratio | (N3-N2) EEG-EGG PAC | 56 | 0.196 | 0.1470 | 0.4843 |
| % Time in Wake | N2 coupled $\kappa$ | 56 | 0.175 | 0.1970 | 0.5949 |
| Sleep to Wake Trans/hr | N2 uncoupled $\kappa$ | 56 | 0.173 | 0.2018 | 0.5949 |
| Delta/Sigma power ratio | EGG ISO var | 56 | 0.162 | 0.2322 | 0.6064 |
| Delta/Sigma power ratio | N3 uncoupled $\kappa$ | 56 | 0.162 | 0.2343 | 0.6064 |
| Delta/Sigma power ratio | N2 coupled $\kappa$ | 56 | −0.160 | 0.2382 | 0.6064 |
| % Time in N1 | N2 uncoupled $\kappa$ | 56 | 0.151 | 0.2673 | 0.6273 |
| Subjective Sleep Quality | (N3-N2) EEG-EGG PAC | 56 | 0.150 | 0.2711 | 0.6273 |
| % Time in N1 | N3 coupled $\kappa$ | 54 | 0.146 | 0.2907 | 0.6273 |
